# Plasmid-encoded insertion sequences promote rapid adaptation in clinical enterobacteria

**DOI:** 10.1101/2024.03.01.582297

**Authors:** Jorge Sastre-Dominguez, Javier DelaFuente, Laura Toribio-Celestino, Cristina Herencias, Pedro Herrador-Gómez, Coloma Costas, Marta Hernández-García, Rafael Cantón, Jerónimo Rodríguez-Beltrán, Alfonso Santos-Lopez, Alvaro San Millan

## Abstract

Plasmids are extrachromosomal genetic elements commonly found in bacteria. Plasmids are known to fuel bacterial evolution through horizontal gene transfer (HGT), but recent analyses indicate that they can also promote intragenomic adaptations. However, the role of plasmids as catalysts of bacterial evolution beyond HGT remains poorly explored. In this study, we investigate the impact of a widespread conjugative plasmid, pOXA-48, on the evolution of various multidrug-resistant clinical enterobacteria. Combining experimental and within-patient evolution analyses, we unveil that plasmid pOXA-48 promotes bacterial evolution through the transposition of plasmid-encoded insertion sequence 1 (IS1) elements. Specifically, IS1-mediated gene inactivations expedite the adaptation rate of clinical strains *in vitro* and foster within-patient adaptation in the gut microbiota. We decipher the mechanism underlying the plasmid-mediated surge in IS1 transposition, revealing a negative feedback loop regulated by the genomic copy number of IS1. Given the overrepresentation of IS elements in bacterial plasmids, our findings propose that plasmid-mediated IS transposition represents a crucial mechanism for swift bacterial adaptation.

## Introduction

Horizontal Gene Transfer (HGT) plays a major role in bacterial evolution. The lateral acquisition of new traits increases genomic diversity and facilitates bacterial adaptation to new environments^1,2^. Conjugative plasmids are key drivers of HGT by spreading accessory genes between diverse bacterial lineages through direct cell-to-cell connections^3,4^. One of the most important and alarming consequences of plasmid-mediated HGT is the global spread of antimicrobial resistance^5,6^ (AMR). The central role of plasmids in AMR spread is highlighted by the wide variety of antibiotics to which they confer resistance, including last-resort antibiotics^7^. AMR acquisition is especially worrying in clinical bacteria, which produce infections difficult to treat and increase mortality rates of hospitalized patients^8^.

Recent studies have revealed that plasmids play an important role in bacterial evolution beyond HGT. For example, plasmids can catalyze evolution by increasing mutation rates of plasmid-encoded genes^9^, maintaining allelic diversity^10^ or promoting phenotypic noise^11–13^. Another important but neglected consequence of plasmid acquisition beyond HGT are intragenomic interactions^14,15^. Plasmids can impact the expression and evolution of loci encoded elsewhere in the bacterial genome. For example, plasmids can produce massive alterations in chromosomal transcriptional profiles^16,17^. In some cases, these alterations can be mediated by plasmid-encoded regulators, which manipulate chromosomal expression to facilitate plasmid maintenance^18^. Moreover, plasmids can directly affect chromosomal evolution through transient hypermutagenesis induced by plasmid-encoded error-prone DNA polymerases^19^. Additionally, plasmids can affect the expression of other mobile genetic elements^18,20^ (MGEs) present in the bacterial genome, such as plasmids or insertion sequences (ISs), and this can in turn promote intragenomic rearrangements^14,17,21,22^. All these intragenomic interactions can affect the evolutionary trajectories of plasmid-carrying bacteria.

ISs are the simplest and smallest transposable elements (TEs), consisting of short DNA segments that encode the enzymes necessary for their own transposition^23,24^. Several examples across the tree of life demonstrate the importance of TEs in evolution^25,26^, from their discovery by Barbara McClintock studying corn kernel colouration^27^, to the emergence of human pathogens^25^, or even the evolution of mammalian lactation and pregnancy^28^. In prokaryotes, ISs can both inactivate genes by disrupting them^29^ or activate/modify^30^ the expression of adjacent genes^31^. IS elements can therefore promote bacterial evolution^22,31,32^, most notably in early stages of adaptation to a new environment^33,34^, as their activity has been suggested to be detrimental in the long term^34^. ISs also provide plasmids with adaptive traits, for example, by flanking regions carrying beneficial genes and conforming composite transposons that can move between plasmids^35^ or between plasmids and the chromosome^36,37^. This mechanism is especially important in AMR spread, as antibiotics can select for the transposition of resistance genes from the chromosome to plasmids^38^.

Accordingly, genomic data suggest a significant overrepresentation of IS elements in conjugative plasmids compared to bacterial chromosomes^23^, as well as a complex network of plasmid/IS interactions shaping the dissemination of AMR^39^. Despite the evident association between ISs and conjugative plasmids^39^, their joint contribution to bacterial evolution beyond its role in AMR dissemination has received little attention.

Importantly, our understanding of plasmids’ effects on the evolution of bacterial pathogens is constrained by two other important limitations. First, most of the works in the field study plasmid-bacteria associations with little or no clinical relevance. Second, previous works have primarily focused on understanding the compensatory evolution of plasmid-associated fitness costs. Nonetheless, recent evidence suggests that plasmids typically impose mild or negligible fitness costs on the majority of their natural bacterial hosts^21,40,41^. To overcome these limitations, in this study we explored the impact of a conjugative AMR plasmid of great clinical relevance, pOXA-48, on the evolution of a wide range of multidrug-resistant clinical enterobacteria^42^. Notably, pOXA-48 produces mild fitness effects in these strains. We combined experimental evolution, whole genome sequencing and transcriptomics of clinical strains with analyses of genomic data obtained from temporal series of isolates recovered from the gut microbiota of hospitalized patients. Our results showed that plasmid-encoded ISs promote rapid adaptation through transposition into the bacterial genome, revealing a new mechanism for plasmid-mediated adaptation beyond HGT.

## Results

### Experimental evolution of pOXA-48-carrying clinical strains

pOXA-48 is a world-wide distributed conjugative plasmid belonging to the plasmid taxonomic unit L/M. This plasmid encodes the OXA-48 carbapenemase and is usually associated with clinical enterobacteria, especially with *Klebsiella pneumoniae* high-risk clones (e.g. ST11 or ST307), which commonly produce nosocomial outbreaks^43^. In recent years we have characterized hundreds of pOXA-48-carrying clinical enterobacteria recovered from patients admitted at the Ramon y Cajal University Hospital in Madrid (R-GNOSIS collection^44^). To study the effect of pOXA-48 in the adaptation of clinical enterobacteria, we followed an experimental evolution (EE) approach with a subset of 13 clinical isolates belonging to three different species: *K. pneumoniae* (n=7), *E. coli* (n=4) and *C. freundii* (n=2) (Methods; Figure 1A and Suppl. Table 1). These clinical enterobacteria are representative of the R-GNOSIS collection both in terms of phylogeny and of pOXA-48 fitness effects in the absence of antibiotics^40,41^ (Figure 1A). Specifically, only one of the strains showed a significant fitness reduction associated with pOXA-48 carriage (Figure 1B). Note that since pOXA-48 produces mild fitness effects in most of these clinical enterobacteria, we could analyze the effects of pOXA-48 on bacterial evolution beyond compensatory adaptation.

**Figure 1.**
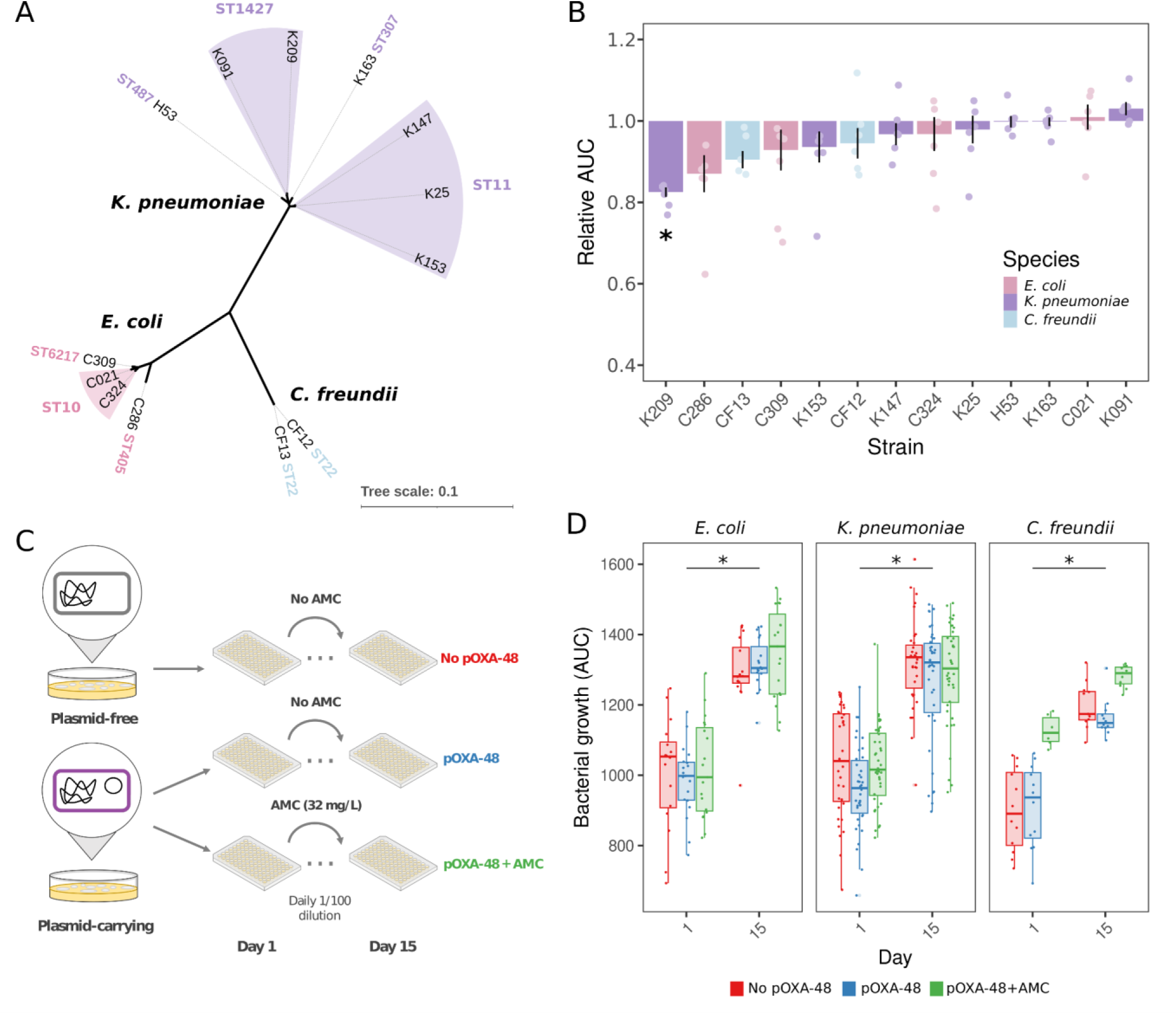
Experimental evolution of clinical enterobacteria. **A)** Strains used in the EE assay. Phylogenetic tree of the strains included in this study. We selected strains from three different species, including the most representative STs from a collection of extended spectrum ß-lactamase (ESBL) or carbapenemase-producing enterobacteria isolated from hospitalized patients (R-GNOSIS collection). **B)** Distribution of fitness effects of pOXA-48 in the different strains included in this study in the absence of antibiotics. We represent the relative Area Under the growth Curve (AUC) of pOXA-48-carrying clones compared to the isogenic pOXA-48-free clones to estimate the plasmid fitness effects. Asterisks indicate those strains in which the plasmid imposed a significant cost (paired t-tests after Bonferroni correction; p < 0.05). **C)** Experimental design of the EE assay performed in this work for 13 bacterial strains. We propagated bacterial populations with and without pOXA-48 in LB for 15 days (∼100 generations, 1:100 daily dilution). We also propagated pOXA-48 carrying bacteria in LB with subinhibitory concentrations of amoxicillin with clavulanic acid (AMC; lower row). **D:** Bacterial growth (AUC) at the beginning (Day 1) and end (Day 15) of the EE (OD600 a.u.·min). We observed a significant increase in the AUC for all of the species in the three conditions from day 1 to 15. The presence of pOXA-48 caused no effect in the increase of AUC.

We propagated isogenic versions of each strain with and without pOXA-48 in LB medium in absence of antibiotics for 100 generations (i.e. 15 days with a daily dilution of 1:100; 6 biological replicates per strain, see Methods and Fig. 1C). In parallel, to examine whether antibiotics affect the evolution of pOXA-48-carrying strains, we also propagated these strains in LB with amoxicillin and clavulanic acid (AMC, 32 mg/L amoxicillin + 6.4 mg/L clavulanic acid), an antibiotic combination to which OXA-48 confers resistance (Fig 1C). To estimate changes in fitness over the EE, we recorded 24-hours bacterial growth curves every alternative day for all the evolving populations (Methods; Suppl. Fig. 1). We observed a significant increase in bacterial growth in the three experimental conditions at the end of the EE (Kruskal-Wallis chi-squared = 229.13, df = 5, p < 0.001). We did not observe a clear impact in fitness changes due to the presence of pOXA-48 (Wilcoxon rank-sum test after significant Kruskal-Wallis test, W = 17507, p = 0.774), and only a marginally positive effect due to the antibiotic pressure (Wilcoxon rank-sum test after significant Kruskal-Wallis test; W = 15663, p = 0.047, Fig. 1D). Overall, these results showed that all populations increased their fitness over the EE, and that pOXA-48 did not significantly affect the adaptation of the clinical strains at the phenotypic level. At the end of the experiment, we confirmed that pOXA-48 remained stable in the evolved populations (see Methods).

### pOXA-48 is associated with IS1 transposition during evolution

To investigate the molecular basis of adaptation during EE, we sequenced the genomes of the evolved populations. We selected the three replicate populations per strain and condition that showed the highest growth at the end of the experiment (n=117). We performed whole-genome sequencing of the evolved populations (short-read Illumina technology at depth > 200x), and compared them with the closed genome of their respective ancestor for each strain (see Methods). With this approach we obtained an overview of the mutations accumulated during the experiment, capturing multiple events at diverse frequencies (Suppl. Table 2, Suppl. Fig. 2). We divided the mutations among SNPs, short insertions/deletions (INDELs), and New Junction (NJ) events. NJ are sequences that are detected jointly in the evolved strains but are in distant sites in the ancestors (insertions/deletions larger than read length^45^). We detected two hypermutator lines at the end of the experiment (affecting *mutS* in a replicate of *E. coli* C324 without pOXA-48 and *mutH* in a replicate with pOXA-48 and AMC of *K. pneumoniae* K147), which were kept out of the final analyses. The spectrum of SNPs at day 15 showed a strong signature of positive selection for the three species in all the conditions (Non-synonymous/Synonymous >> 1.0, Suppl. Table 3), as well as a notable enrichment in intergenic mutations (Fig. 2; Suppl. Table 3). pOXA-48 did not affect the frequency of SNPs in the populations of *E. coli* strains (Chi-squared = 4.1391, df = 6, p = 0.658) but it did in *C. freundii* and *K. pneumoniae* ones, which normally showed less SNPs in presence of the plasmid (Chi-squared = 27.703, df = 6, p < 0.001).

**Figure 2.**
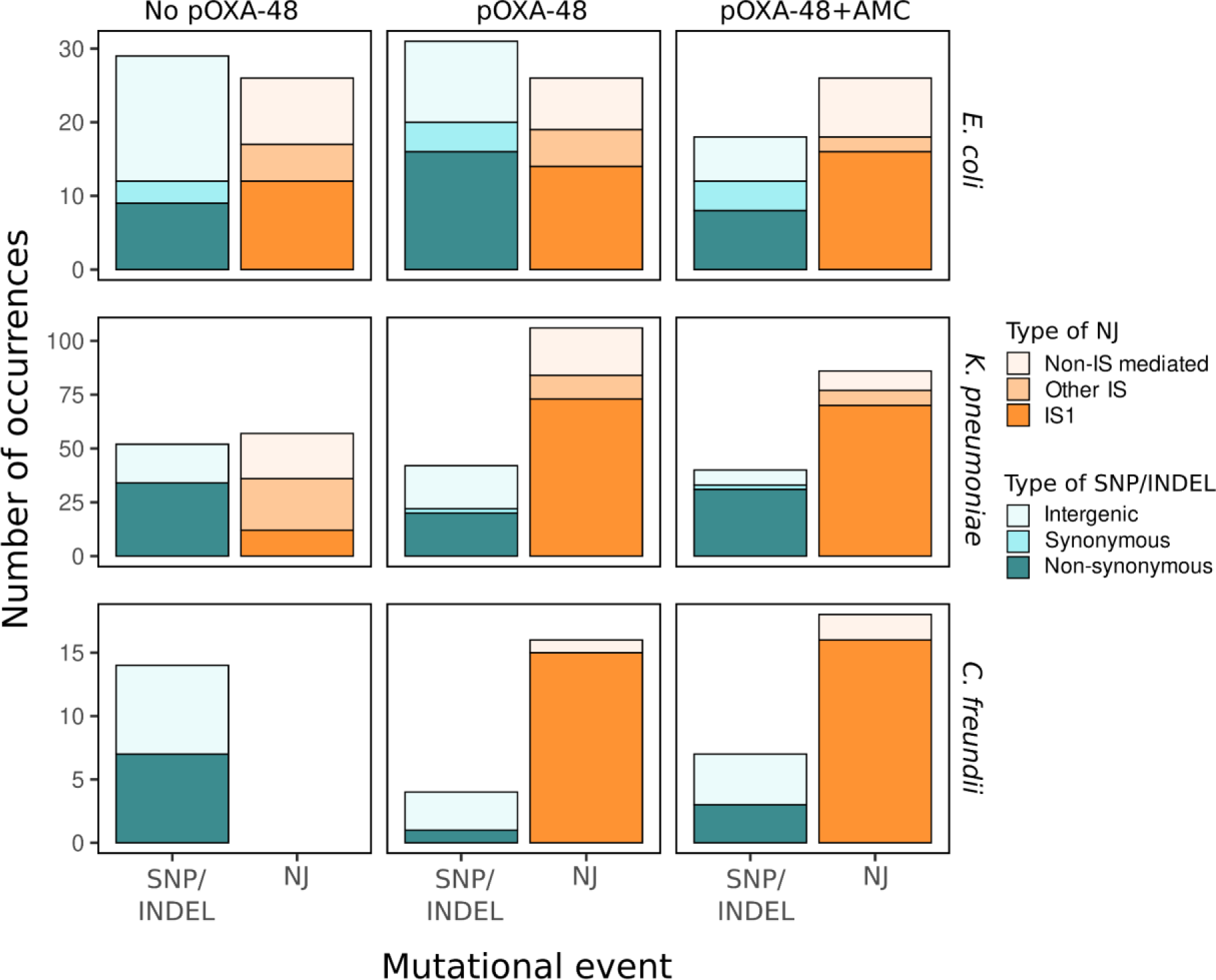
Overview of the mutational profiles in the evolved populations. Summary of the genetic changes in the evolved populations in the different conditions of the experiment. Barplots represent the total number of occurrences of each type of event in all propagated populations depending on the experimental condition and species. We summarized the different events as the addition of each type of mutation for all the strains of each species in each condition (note that we used 7 *K. pneumoniae*, 4 *E. coli* and 2 *C. freundii* strains). Events were divided depending on their nature (i.e. if they were Single Nucleotide Polymorphisms (SNPs)/small INDELs or New Junctions (NJ)), and their type (Intergenic, Non-Synonymous or Synonymous SNPs/INDELs; IS1-mediated, Non-IS-mediated, or mediated by other ISs NJs). We classified INDELs as Non-synonymous if they were found in coding regions or Intergenic for those in intergenic regions. We noticed a considerable increase in IS1-mediated NJ events in presence of the plasmid, both in *K. pneumoniae* and *C. freundii*, but not in *E. coli* (Suppl. Fig. 3).

NJ events were common during the EE assay (Fig. 2, Suppl. Fig 3). Overall, the majority of NJ events reported were mediated by the IS1 (69.5% in *E. coli*; 79.92% in *K. pneumoniae*; 97.14% in *C. freundii*). Interestingly, pOXA-48 carries two copies of the IS1, and the different ancestral strains carried a variable number of additional IS1 copies in their genomes, ranging from 0 to 40 (Suppl. Table 1). In our sample, *E. coli* strains encoded a significantly higher number of IS1 genomic copies (average = 29.5, sd = 10.15), compared to *K. pneumoniae* (average = 3.7, sd = 1.6) and *C. freundii* strains (average = 1.5, sd = 0.7), (Kruskal-Wallis chi-squared = 9.0712, df = 2, p = 0.011). In the *E. coli* populations, the overall mutational profiles were conserved across all the experimental conditions, showing a similar number of IS-mediated NJ in the different treatments (two-sided Kruskal-Wallis test, chi-squared = 0.33163, df = 2, p = 0.8472). However, in *K. pneumoniae* and *C. freundii,* we observed a considerable increase in IS-mediated NJ events in pOXA-48-bearing populations compared to the pOXA-free ones (Fig. 2, two-sided Kruskal-Wallis test; *K. pneumoniae* chi-squared = 18.551, df = 2, p < 0.001; *C. freundii* chi-squared = 9.9479, df = 2, p = 0.007; Suppl. Fig. 3, Suppl. Fig. 4).

### pOXA-48-encoded IS1 promotes adaptation in K. pneumoniae and C. freundii

Our results suggested that the presence of pOXA-48 may be associated with a higher IS1 transposition rate in *K. pneumoniae* and *C. freundii*. To investigate if IS1 played an important role driving bacterial adaptation during the EE, we analyzed the targets that showed parallel evolution at the end of the experiment across all the different conditions. Genes mutated in parallel across conditions and populations are a strong indication of targets under positive selection^46,47^ (Fig. 3AB). Genomic analyses revealed multiple parallel targets across different replicates, strains and even species (Fig. 3AB; Suppl. Fig. 5-17; full table of mutations in Suppl. Table 2). In *E. coli* and *C. freundii*, the main targets were the fimbrial operon (*fimABCDE*) and a stress response operon^48^ (*rpoS*, *nlpD*, Fig. 3C). In *K. pneumoniae*, the fimbrial operon was also highly targeted, as well as the capsule operon (*mkrJ*, *wbaP*, *wcaJ* and diverse glycosyltransferase-coding genes, Fig. 3D). The inactivation of these targets has been previously described during EE assays in enterobacteria. For instance, the most common adaptation in *K. pneumoniae* in our experiment, capsule inactivation, has been described to enhance fitness *in vitro*^49^. In our experiment, these *loci* were affected both in absence and in presence of pOXA-48 and antibiotics, indicating that they promoted general adaptation to the experimental conditions. These targets were mostly altered via SNPs or small INDELs in *E. coli* populations and in pOXA-48-free *K. pneumoniae and C. freundii* populations. However, in pOXA-48 carrying *K. pneumoniae* and *C. freundii* populations, the targets were mainly disrupted by IS1 insertions (Fig 3D and Suppl. Fig. 5-17). To confirm the results obtained from the WGS of the whole populations, we sequenced the genomes of clones from each evolved *E. coli* and *K. pneumoniae* population, combining short and long read sequencing technologies (n=14; 1-3 clones per population). The closed genomes generated allowed us to confirm the movements of the IS1 observed in the population analyses (Suppl. Table 4).

**Figure 3.**
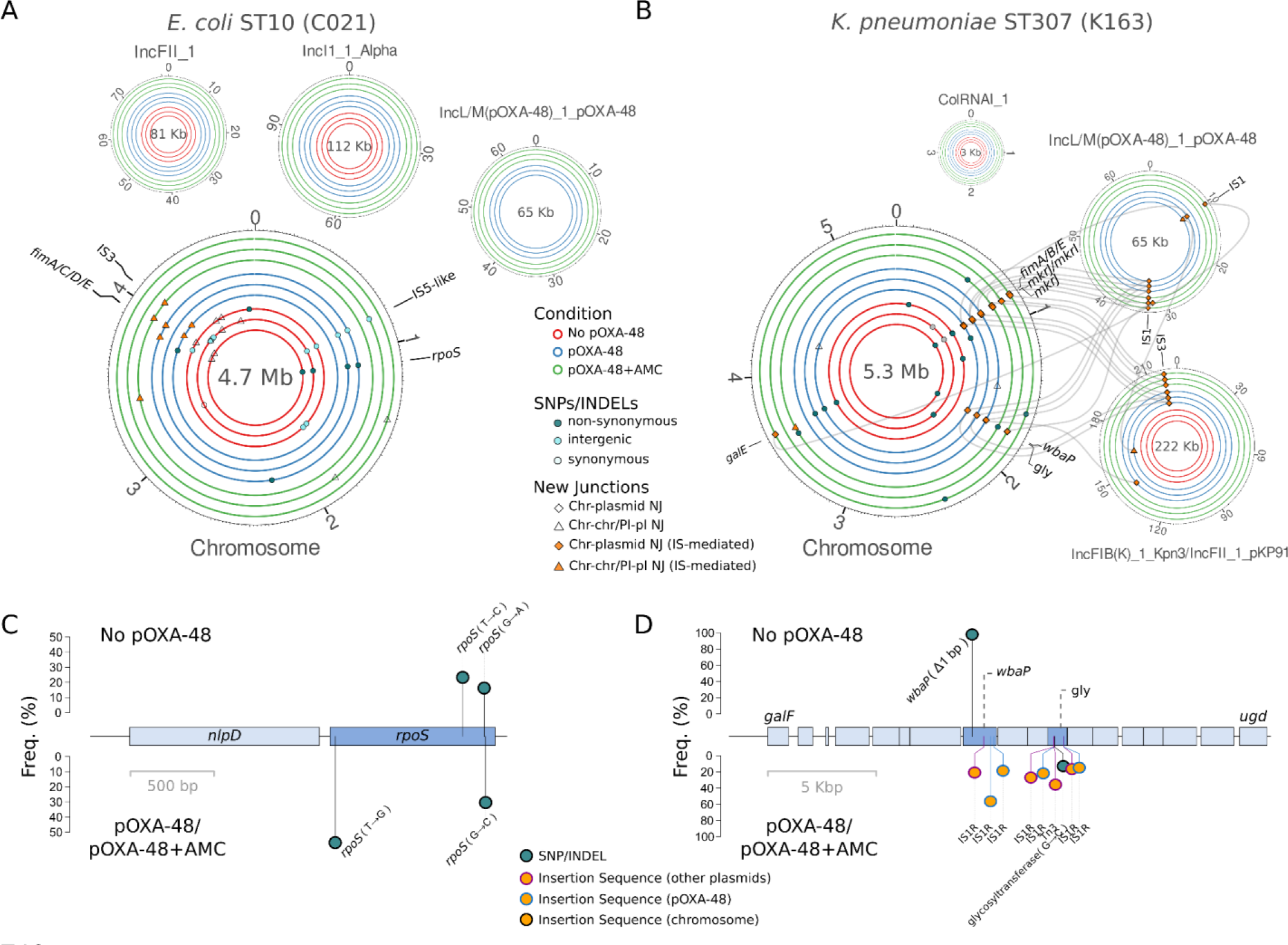
Adaptive targets in the species under study. **A, B)** Plots showing the mutations in the genomes of an *E. coli* (A), and a *K. pneumoniae* (B) strains selected as examples of each species (see Suppl. Figures 5-17 for all the strains). From inside to outside, circles indicate the different replicates of the evolved bacterial populations without pOXA-48 (red), carrying pOXA-48 (blue) and carrying pOXA-48 in presence of AMC (green). Units represent megabases (Mb) in chromosome replicons and kilobases (Kb) in plasmid replicons. Parallel evolution targets are labeled in each circa plot. The different types of SNPs/INDELs are represented by dots, whereas NJ events are depicted by shapes, filled in orange in the case of IS-mediated NJs. IS rearrangements which could be tracked (i.e., confirmed by genomic data, and/or by long-read sequencing) are shown as lines connecting the IS element and its target. Note that as the data of fully evolved populations is shown, it is possible to encounter multiple mutations or insertions at intermediate frequencies within the same gene. **C, D)** Operons parallelly targeted during evolution in our experiment. In the case of *E. coli* (C) and *C. freundii*, the *nlpD-rpoS* operon was targeted in multiple strains (Suppl. Fig. 5-10). In almost all of the *K. pneumoniae* strains, the capsule operon (D) was mutated at the end of the evolution (Suppl. Fig. 11-17). The top part of the plots show mutations occurred during the EE in the replicates propagated in absence of the plasmid, whereas the bottom part includes the events that happened in pOXA-48 carrying replicates both with and without AMC. Freq. indicates the frequency of the mutations/insertions in the population in percentage. Names of the genes are shown. Function of those genes not named by PGAP are also indicated: gly: glycosyltransferase.

As we mentioned before, the strains under study carried a variable number of IS1 elements in their genomes. It was therefore difficult to determine whether the IS1 movements recorded during EE had originated from pOXA-48 or from the remaining IS1 copies. However, in a subset of the strains tested, we could confirm the transposition of pOXA-48-encoded IS1 due to differences in its genetic sequence compared to the remaining IS1 elements in the genome (3 *K. pneumoniae* and 1 *C. freundii,* Fig. 3B and Suppl. Fig 10, 13 and 17; see Methods). This result suggested that pOXA-48-encoded IS1s were responsible for most of the IS1 transposition events observed during EE in these strains (Suppl. Fig. 10, 13 and 17).

### IS1 genomic copy number determines the frequency of transposition

We decided to investigate the differences in IS1 transposition frequencies associated with the presence of pOXA-48 in the species under study. The IS1 encodes two partly overlapping open reading frames, *insA* and *insB*^50^. The active transposase, InsAB’, is synthesized through a low-frequency translational frameshift in the overlapping region^50^. The InsA protein, on the other hand, represses the expression of the IS1-encoded genes^51^. We hypothesized that the InsA proteins present in the strains could be repressing the transposition of the incoming pOXA-48 IS1 elements. Supporting this idea, we found a negative correlation between the pre-existing number of IS1 copies in the genome of the strains and the number of IS1 transposition events during EE in pOXA-48-carrying strains (n=13; Fig. 4A; Pearson correlation R = -0.57, p = 0.028). To further investigate this hypothesis, we analyzed RNA-Seq data from a subset of isogenic strain pairs with and without pOXA-48 belonging to the R-GNOSIS collection (Suppl. Table 1). We also found a negative correlation between the number of pre-existing IS1 copies and the increase in *insAB* transcription levels in presence of pOXA-48 (n=10; Fig. 4B; Spearman correlation Rρ = -0.75, p = 0.012).

**Figure 4.**
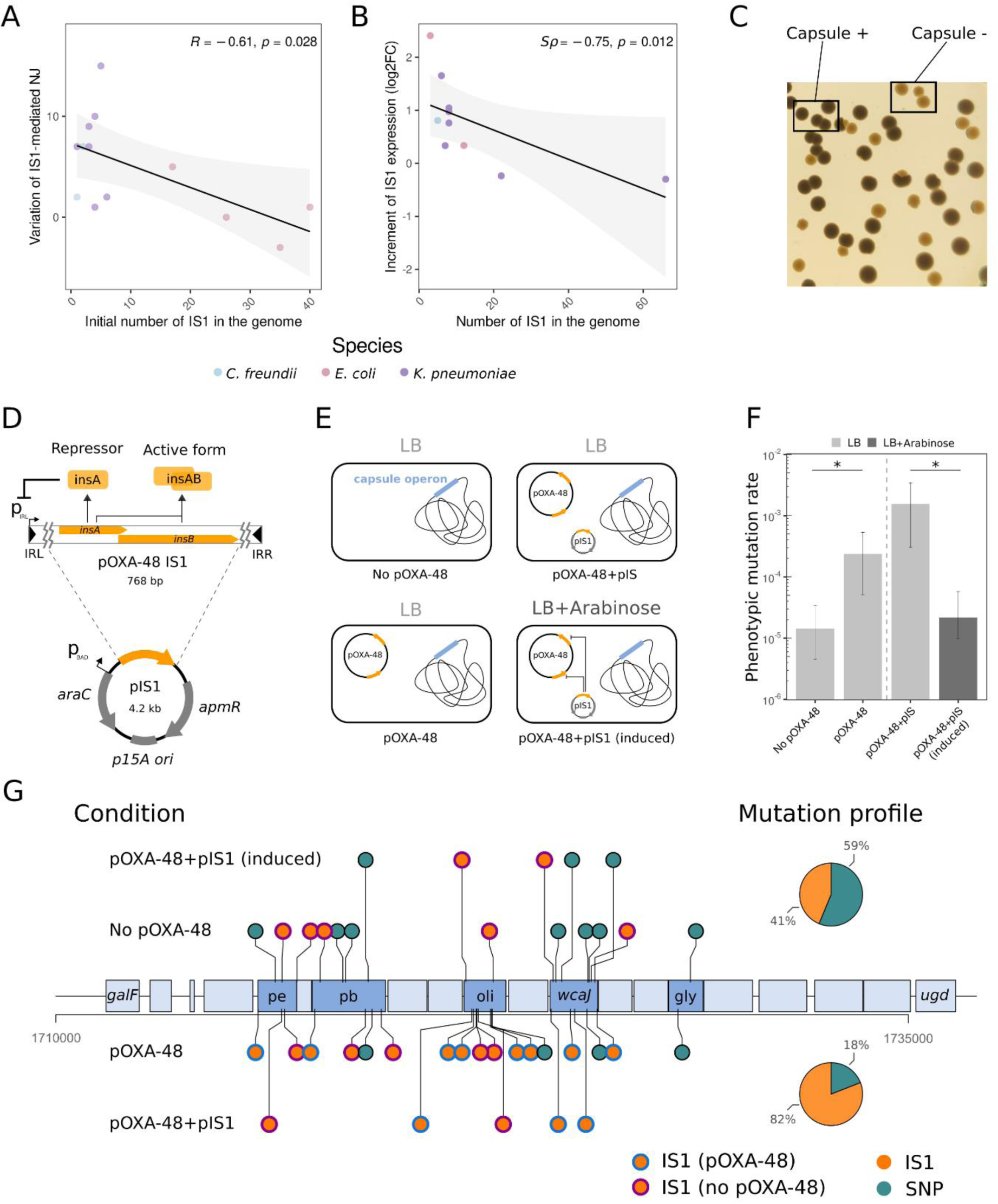
Regulation of IS1 transposition in pOXA-48-carrying strains. **A)** Pearson correlation between the number of IS1 copies in the genomes of the ancestral strains at the beginning of the experimental evolution and the increment in IS1 copies during the experiment. **(B)** Spearman correlation between the change of expression of IS1 elements after pOXA-48 acquisition and the number of IS1 copies in the genome of multiple strains from the R-GNOSIS collection (note that some of these strains are the same as those in the EE: n=5). **C)** *K. pneumoniae* (K25 strain) colonies spotted on an LB agar plate either capsulated (darker colonies) or capsule-free mutants (lighter colonies). **D)** Schematic representation of plasmid pIS1. We cloned the IS1 element of pOXA-48 keeping out the inverted repeats (IRL and IRR) that contain its promoter. The arabinose-inducible promoter P_BAD_ controls the expression of the *insAB* genes. pIS1 also encodes an apramycin resistance gene for selection. **E)** Experimental design including the different K25 versions generated for the fluctuation assay to test loss-of-capsule mutation rate. **F)** Loss-of-capsule phenotypic mutation rate in the different conditions tested. Error bars indicate the 95% confidence interval. Asterisks denote significant differences (p < 0.05) between the conditions tested according to likelihood ratio tests. **G)** Mutations detected at the end of the fluctuation assay in capsule-free clones from the different conditions. Mutations accumulated in absence of pOXA-48, or under conditions of ins*AB* expression from pIS1 are represented in the upper side of the panel. Mutations accumulated in the presence of pOXA-48, or pOXA-48 + pIS1 without induction are represented in the lower part of the panel. Line color of IS1 insertions indicate the origin of the sequence (pOXA-48 or no pOXA-48). The name of the genes is indicated inside of their coding region. The function is shown for those capsule genes not named by PGAP: pe: polysaccharide export protein; pb: polysaccharide biosynthesis tyrosine autokinase; oli: O-antigen ligase; gly: glycosyltransferase.

To experimentally investigate the effect of IS1 copy number on the frequency of IS1 transposition, we designed an experiment to quantify the mutation rate leading to capsule loss in *K. pneumoniae*. Capsule loss mediated by gene inactivations was one of the main adaptations observed during the EE in *K. pneumoniae* (Fig. 3 and Suppl. Fig. 11-17) and produced a conspicuous phenotype easy to spot on the agar plates (i.e. transparent colonies; Fig. 4C). To modify IS1 dosage, we cloned the *insAB* ORFs under the control of an arabinose-inducible promoter located in a small multicopy plasmid, pIS1 (Fig 4B; Suppl. Fig. 18; Methods). We then selected one of the *K. pneumoniae* strains used in the EE assay that carries a low number of IS1 elements in the genome (K25, n=1). We introduced pIS1 both in the pOXA-48-carrying and pOXA-48-free K25 strains (Fig. 4E). Next, we performed fluctuation assays with and without *insAB* induction and calculated the mutation rate leading to the loss-of-capsule phenotype in each treatment^52^ (Fig 4F, see Suppl. Fig. 18 for all the controls). As expected from the EE results, we detected a significantly higher number of phenotypic capsule-free mutants in the pOXA-48-carrying strain than in the pOXA-48-free one (Fig. 4F, likelihood ratio test, p < 0.05). Interestingly, the increase in capsule-free mutants was completely reverted in the pOXA-48-carrying strain when *insAB* expression was induced, suggesting that a high dosage of InsA reduces IS1 transposition (Fig. 4F; likelihood ratio test, p > 0.05). Genome sequencing of a subset of clones from these assays confirmed that capsule inactivation was mostly mediated by IS1 in the pOXA-48-carrying strain and in the pOXA-48-carrying strain with a non induced pIS1 (82%, n=23; Fig. 4G), but not in the pOXA-48-free strain or in the pOXA-48-carrying strain with *insAB* overexpression (41%, n=17; Fig. 4G).

Importantly, the capsule genes mutated in the fluctuation assay were the same as those mutated during the EE assay in this strain (Suppl. Figure 14; Suppl. Table 5). Taken together, our results confirmed that the number of IS1 genomic copies modulates pOXA-48-associated IS1 transposition frequencies, explaining the different mutational profiles observed for each species in the EE.

Finally, we performed competition experiments between a subset of the mutants obtained in the fluctuation assay (which carried no other mutation in the genome) and the capsulated strain. Results confirmed that capsule loss provides a fitness advantage under our experimental conditions regardless of the pOXA-48 presence (Suppl. Fig. 18 C).

### pOXA-48-encoded IS1 promotes within-patient evolution in clinical enterobacteria

One important limitation associated with EE assays is the difficulty of extrapolating the evolutionary dynamics observed beyond the specific experimental conditions tested^53,54^. To overcome this limitation, we decided to study IS1 dynamics directly in sequential clinical enterobacteria colonizing the gut microbiota of hospitalized patients. In our previous studies with the R-GNOSIS collection, we described multiple temporal series of pOXA-48-carrying enterobacteria recovered from patients^55,56^. To track within-patient IS1 movements, we screened our collection for patients where the same pOXA-48-carrying lineage was isolated more than once during hospitalization. We classified isolates as belonging to the same lineage if they differed in less than 7 SNPs across the entire genome^57^. We detected 27 patients fulfilling this condition, from which 69 isolates belonging to 30 different lineages were recovered: *E. coli* = 10, *K. pneumoniae* = 19, *C. freundii* = 1 (Suppl. Table 1, see Methods). As observed before, in this new set of strains *E. coli* also encoded a significantly higher number of IS1 genomic copies compared to *K. pneumoniae* and *C. freundii* (Fig. 5A; Kruskal-Wallis test p < 0.001)

**Figure 5.**
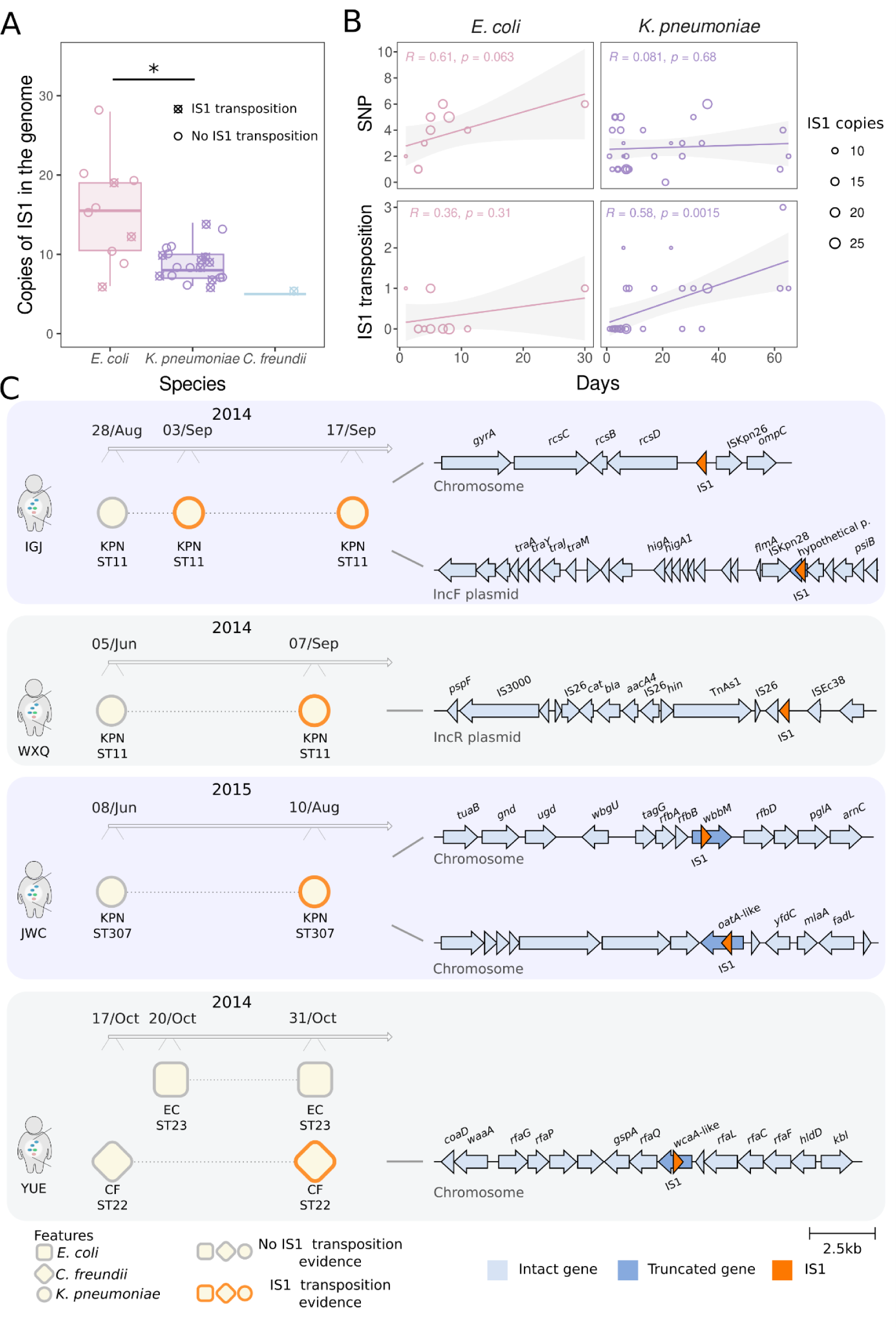
Within-patient evolution of pOXA-48-carrying enterobacteria. **A)** Number of genomic IS1 copies in the strains analyzed. Number of IS1 and IS1-like elements annotated in the genomes of each of the strains. We detected a significantly higher number of IS1 elements in the genomes of the *E. coli* strains analyzed. Crossed dots denote strains in which IS1 movements were detected. Asterisk denotes significant differences (Kruskal-Wallis test p < 0.05) between species. Due to the low sample size of *C. freundii* isolates (n=1), we could not perform a comparison including this group. **B)** Within-patient mutation dynamics. Representation of the number of IS1 transposition events or SNPs/INDELs as a function of time (days) between isolations for the different *E. coli* and *K. pneumoniae* lineages under study. The diameter of the points is proportional to the IS1 genomic copy number of the strains. **C)** Targets of pOXA-48-mediated IS1 transposition in the gut of hospitalized patients. Left side of the panel shows the timelines of the isolation of pOXA-48-carrying enterobacteria from patients IGJ, WXQ, JWC and YUE. Isolates features such as species, ST and IS1 transposition evidence in the lineages are detailed. Isolation dates are also indicated in the timeline. Right side of the panel shows the regions of the genome in which the IS1 element inserted in each of the lineages during *in vivo* evolution. The replicon in which we detected the insertion is indicated. In some of the lineages we could detect more than 1 transposition event (patients IGJ and JWC). Genes disrupted by IS1 elements are highlighted in darker blue. The names of the genes annotated by PGAP are shown on top.

To analyze IS1 movements in these lineages, we combined short- and long-read DNA sequencing. We first compared the genomes of the different isolates from each lineage to determine the genetic changes accumulated over time (Fig. 5B). We detected IS1 transposition events in 3/10 *E. coli*, 9/19 *K. pneumoniae* and 1/1 *C. freundii* lineages from 13 different patients (Suppl. Table 6). Again, a subset of the strains analyzed showed distinguishable pOXA-48-encoded IS1 elements when compared to the remaining IS1 elements in the genome (n=11; 3 *E. coli*, 7 *K. pneumoniae* and 1 *C. freundii*). Among these, we detected pOXA-48-mediated IS1 transposition events in 3 *K. pneumoniae* lineages and 1 *C. freundii* lineage, but not in *E. coli* (Figure 5C). Taken together, these results suggest that pOXA-48 promotes IS1 transposition in the gut microbiota of patients, especially when the genomic IS1 copy number is low (Fig. 5A-B). To test this idea, we built a generalized additive model interrogating the effect of the initial number of IS1 genomic copies in the within patient IS1 transposition rate (see Methods). Results revealed that the number of IS1 copies in the genome, but not strain species, negatively affected IS1 transpositions frequency (IS1 copies: F = 3.28; R-squared = 0.203, p = 0.032; Deviance explained = 29.7%; species: Intercept t = - 0.704, p = 0.486; *E. coli*, t = 1.73, p = 0.093; *K. pneumoniae,* t = 0.1654, p = 0.437; Suppl. Fig. 19).

Finally, we investigated the within-patient targets of IS1 transposition. We analyzed the 4 patients where we could specifically track the IS1 transposition from pOXA-48 to different targets in the genomes of 3 *K. pneumoniae* and 1 *C. freundii* lineages. Interestingly, in three out of four patients, these targets included loci associated with the lipopolysaccharide (LPS) or O-antigen composition, which are well known adaptive targets during within-patient evolution of bacterial pathogens^46,58^ (i.e. pathoadaptive mutations; Fig. 5C). In patient IGJ, the target was the Regulator of Capsule Synthesis system, involved in stress response and virulence regulation^59^. In patient JWC the gene mutated was *wbbM* in the *rfb* locus, which inactivation has been previously associated with O-serotype modification and LPS defects^60,61^. Interestingly, an analysis of *K. pneumoniae* genomes (n > 8000) found the *rfb* locus to be commonly disrupted by a variety of IS elements^60^. The other target found in this patient was an *oatA*-like (∼30% homology) acyltransferase. Mutants in this gene have been associated with immune escape in other species^62^. Finally, in patient YUE, the target of transposition was a *wcaA*-like (∼30% homology glycosyltransferase encoded in the *rfa* locus), which is involved in the core structure of the LPS^63^. Altogether, these results suggest that pOXA-48 may promote within-patient pathoadaptation in *K. pneumoniae* high-risk clones, such as those belonging to ST11 or ST307.

## Discussion

In this study, we explored the impact of an AMR plasmid of great relevance, pOXA-48, on the evolution of clinical enterobacteria beyond HGT (Fig. 1). Informed by recent advances in the field, we designed our experimental system to be able to explore pOXA-48 general effects in bacterial adaptation, instead of focusing on compensatory evolution of plasmid-associated fitness costs. Results revealed that plasmid pOXA-48 is associated with an increase in IS1 transposition rate that promotes bacterial adaptation through gene inactivation *in vitro* (Figs. 2-3). In addition, results from clinical enterobacteria recovered from the gut microbiota of patients strongly suggested that pOXA-48-mediated IS1 transposition also promotes bacterial adaptation *in vivo* (Fig. 5). IS1 transposition is regulated by a negative feed-back loop, so the pre-existing IS1 copy number in the bacterial genome determines the recombination rate of the incoming plasmid-encoded IS1 elements (Fig. 4). These results shed new light into the mechanisms of plasmid-mediated bacterial adaptation.

The key result of our study is that plasmid-mediated IS1 transposition speeds up the rate of adaptation in clinical enterobacteria. Importantly, this effect is dependent on the IS1 genomic copy number. Two implications arise from this observation. First, pOXA-48 can help bacteria to rapidly adapt to strong selective pressures by providing a transient burst of IS1 transposition. In fact, we provide evidence supporting that adaptation of clinical enterobacteria is rapidly driven by IS1 transposition in pOXA-48-carrying strains, both *in vitro* and *in vivo*. This effect could help explain how successful associations between pOXA-48 and high risk bacterial clones emerge in clinical settings, where bacteria face strong selective pressures and changing environments^64,65^. Second, the impact of pOXA-48 on bacterial evolution likely varies significantly from strain to strain. IS1 is present at an average of ∼17 copies per prokaryotic genome^66^, but it is not homogeneously distributed across bacteria. For example, IS1 elements are present at around 3- to 7-fold more copies in *Escherichia* than in *Citrobacter* or *Klebsiella* genomes, respectively^66^. But even at the species levels, there are notable differences in IS1 copies. The *K. pneumoniae* and *C. freundii* strains we used for EE contained, by chance, a considerably lower number of copies than their species average, supporting the idea that these were more prone to an IS1 transposition burst driving adaptation in presence of pOXA-48.

The long-term effects of IS elements on bacterial evolution are poorly understood. There is no obvious explanation of why these elements are maintained within bacterial populations^67^, as they can be beneficial in early stages of adaptation to new environments but they also show deleterious effects on the long run^67,68^. Models based on the fitness effects of ISs claim that both transposition bursts and IS down regulation contribute to the maintenance of IS elements in bacterial populations^67^. Our results indicate that pOXA-48 induces a transient burst of IS1 transposition, especially if the IS1 genomic copy number is low in the recipient bacterium. Importantly, this burst is self-limiting, because once the IS1 copies increase in the genome, transposition is repressed. It is tempting to speculate that this particular transposition tempo may promote persistence of both plasmids and ISs in bacterial populations. Specifically, the initial transposition burst could help resolve conflicts between the incoming plasmids and the resident genome, and the subsequent down-regulation could avoid long-term IS1-associated deleterious effects. Alternatively, IS elements might enjoy the benefits of plasmid-mediated dissemination at the expense of both plasmids and bacterial hosts. Future experiments looking at the outcome of ISs, plasmids, and bacteria at different initial MGE configurations could help to test these hypotheses.

One important limitation of our study is the fact that we only worked with plasmid pOXA-48, so it is difficult to assess if other plasmids promote bacterial adaptation through IS transposition. However, plasmids are generally enriched in IS elements compared to chromosomes^23^, so we argue that this effect could be a general mechanism of plasmid-mediated adaptation in bacteria. In fact, there are previous reports suggesting that gene inactivation through the movement of plasmid-encoded IS, including the IS1 family, promotes AMR evolution in bacteria^69,70^. Future work will help to assess the generality and consequences of plasmid-mediated IS transposition in bacterial evolution. Another limitation of this study is that our experimental work was performed *in vitro*, under classic laboratory conditions. To try to overcome this limitation, we analyzed temporal series of pOXA-48-carrying lineages recovered from patients. Results from this analysis supported our experimental observations, since we detected that IS1 transposition in these lineages anticorrelated with IS1 genomic copy number. Importantly, the targets of IS1 transposition *in vivo* included multiple genes involved in the structure and composition of virulence factors such as the O-antigen or the membrane LPS, which are well described pathoadaptive targets^46,58^. These results suggest that pOXA-48 may not only drive AMR dissemination in high-risk clinical enterobacteria, but it could also promote the evolution of virulence in these strains.

## Methods

### Bacterial strains

We selected the samples included in this work from a collection obtained in the Hospital Universitario Ramón y Cajal as part of an active surveillance-screening programme for detecting patients colonized^44^ by extended-spectrum-beta-lactamases (ESBL) and/or carbapenemase producing Enterobacteriaceae, approved by the Hospital Ethics Committee (Reference 251/13). Samples were isolated and characterized as described before^44^. The bacterial strains used in the EE assay included 7 *Klebsiella pneumoniae* (ST11, ST307, ST487 and ST1427), 4 *Escherichia coli* (ST10, ST131 and ST405) and 2 *Citrobacter freundii* (ST22, see Suppl. Table 1). We considered this subset representative of the R-GNOSIS collection in terms of phylogeny, pOXA-48 variants (Suppl. Fig. 20) and fitness effects. The strains we used to study within-patient evolution of pOXA-48-carrying clinical strains included 10 *E. coli* lineages, 19 *K. pneumoniae* lineages, and 1 *C. freundii* from 27 different patients (see Suppl. Table 1). For the RNA-Seq analysis, we employed 2 *E. coli*, 6 *K. pneumoniae* and 1 *C. freundii* clones (Suppl. Table 1). We performed the phylogenetic analysis using mashtree v.1.2.0 and represented it using the Interactive Tree of Life (iTOL) online tool.

### Experimental evolution

We propagated the above mentioned 13 clinical isolates with and without pOXA-48 obtained as explained in the previous section. To analyze the effect of pOXA-48 during bacterial evolution, we propagated 6 replicate cultures per strain of plasmid-carrying and lacking bacteria. Also, to analyze the effect of antibiotics during evolution, we propagated 6 replicate cultures per strain of plasmid-carrying bacteria in the same way with amoxicillin/clavulanic acid (AMC) at an inhibitory concentration for plasmid-free bacteria, but sub-inhibitory concentration for plasmid-carrying clones (32 mg/L amoxicillin + 6.4 mg/L clavulanic acid). This means that for each strain, at the end of the experiment we had three conditions for the evolved bacteria, named as follows in the manuscript: no pOXA-48 (plasmid-free), pOXA-48 (plasmid-carrying) and pOXA-48+AMC (plasmid-carrying in presence of antibiotic). In total, we propagated 234 independent populations. We used a single colony as the ancestor for each replicate of the plasmid-free replicates. We used 6 independent colonies for the replicates bearing the pOXA-48 propagated with and without antibiotic (*i.e.* each colony started 1 replicate of plasmid-carrying and 1 replicate of plasmid-carrying in presence of antibiotic). For each condition, we propagated 6 replicate cultures in Lysogeny Broth medium (LB, Condalab) for 15 days, at 37°, 250 r.p.m., making serial daily passages of the cultures in 96-well plates (1:100 dilution, 2μl transferred every 24 hours into 200μl of LB, i.e. 6,64 generations per day, which resulted in *ca.* 100 bacterial generations at the end of the EE). We checkerboard-filled the 96-well plates to avoid cross-contamination in the experiment.

To assess the phenotype of clinical enterobacteria during evolution, we analyzed the bacterial growth during the EE assay for all the replicate populations. We measured growth curves every 48 hours. We quantified the optical density (OD600) at 600 nm wavelength every 10 minutes under constant shaking conditions (365 r.p.m. during 8 min. + OD600 measurement during 2 min.) for 24 hours using a plate reader (Synergy HTX Multi-Mode Reader; BioTek Instruments) and using the Gen5 3.11 software. We checked for possible cross-contaminations by plating in HiCromeTM UTI (HiMedia Laboratories, India), which allows us to differentiate between species and by testing the presence of pOXA-48, in pOXA-48 free strains (primers ID-7 to ID-12; Suppl. Table 7). In total we discarded from further analyses 29 replicates with possible cross-contamination events. At the end of the experiment, we estimated plasmid stability by replica plating of *ca.* 50-100 colonies in AMC. The plasmid was 100% stable in all populations but one replicate of C286 which showed a stability of 93%. We stored the propagated populations at days 1, 2, 3, 4, 5, 7, 9, 11, 13 and 15 with 15% glycerol at -80°C.

### pOXA-48 fitness effects

To analyze the fitness effects of pOXA-48 in the different bacterial strains, we measured the bacterial growth as explained in the previous section at day 1 of evolution, and compared the growth of plasmid-carrying *vs* plasmid-free bacteria for each strain. Briefly, we calculated the Area Under the growth Curve (AUC) for each plasmid-carrying and plasmid-free replicate, as it is a parameter that integrates useful growth information and is convenient for the estimation of plasmid fitness effects. We then divided the mean AUC of plasmid-carrying bacteria by the mean AUC of plasmid-free bacteria to determine the fitness effect of pOXA-48 in each strain.

### Whole genome sequencing

#### Short-read Illumina sequencing

We extracted the genomic DNA using the Wizard Genomic DNA Purification Kit (Promega, WI, USA). Illumina sequencing was performed at SeqCoast Genomics (Portsmouth, NH, USA), at the Microbial Genome Sequencing Center (MiGS; Pittsburgh, PA, USA) and at the Wellcome Trust Centre for Human Genetics (WTCHG; Oxford, UK), using the NextSeq 2000 and NovaSeq 6000 platforms. Short reads generated were paired and 150 bp long.

For the EE assay, we performed WGS reaching a coverage of *ca.* 200x on whole populations of the evolutionary ancestor of each strain at day 1 of the EE (1 replicate per condition at day 1, and merged the sequences of the 3 ancestors per strain for the downstream analyses) and on the 3 evolved populations per condition and strain, which showed best growth for each experimental condition per strain (day 15 of experimental evolution). We sequenced the experimental ancestors (day 1) as a control to filter out possible mutations present in the ancestral clones or that appeared during the unfreezing of the strains, and not due to the evolutionary conditions. Finally, we sequenced between one and three clones per evolved population of *E. coli* and *K. pneumoniae* (n=14) used for WGS to confirm the variant calling of structural variants by direct comparison with long-read data (see *Long-read Oxford Nanopore sequencing* subsection). For a simplified summary of EE+WGS methods see Suppl. Fig. 20.

In addition to the EE, we also used short-read Illumina sequencing for the analysis of the fluctuation assays, for studying the evolution of plasmid-encoded IS1 *in vivo* and for closing the reference genomes used in the EE and RNA-Seq analyses (see Methods *Fluctuation assay to test IS1 transposition rate, Analysis of within-patient transposition events* and *Reference genomes* for details).

#### Long-read Oxford Nanopore sequencing

To close the genomes of the ancestral strains of the EE by hybrid assembly, we also sequenced their whole genomes using Oxford Nanopore technologies. Besides, to better determine MGE rearrangements, as well as to confirm structural variants (e.g. the transposition of the IS1 element from pOXA-48), we isolated 14 evolved clones and sequenced them by long read to complement the results obtained from Illumina sequencing. For this, we followed the Nanopore protocol for Native barcoding genomic DNA (with EXP-NBD104, EXP-NBD114, and SQK-LSK109) v. NBE_9065_v109_revAK_14Aug2019. Briefly, we extracted the genomic DNA as explained in the previous section. For the quantification of the extracted DNA we employed a Qubit (Invitrogen™ Qubit Flex Fluorometer, ThermoFisher Scientific). In the DNA repair and end-prep steps we used the NEBNext Ligation Sequencing Kit (Oxford Nanopore Technologies, UK), AMPure XP beads and a Hula mixer (Beckman Coulter™ Agencourt AMPure XP, ThermoFisher Scientific). For the native barcode ligation step we used the EXP-NBD104 and EXP-NBD114 kits (Oxford Nanopore Technologies, UK) as well as the Blunt/TA Ligase Master Mix (New England Biolabs, MA, USA). We performed the adapter ligation and clean-up steps with the NEBNext Quick Ligation Reaction Buffer (New England Biolabs, MA, USA), NEBNext Ligation Sequencing Kit (New England Biolabs, MA, USA) and the Ligation Sequencing Kit (SQK-LSK109, Oxford Nanopore Technologies, Oxford, UK). Finally, for the Priming and loading of the Flow Cell steps we used the Flow Cell Priming Kit (EXP-FLP002, Oxford Nanopore Technologies, Oxford, UK) and sequenced the samples using R9.4.1 Flow Cells (FLO-MIN106D) in a MinION Mk1B.

In addition to the EE, we also used Long-read Oxford Nanopore sequencing for studying the evolution of plasmid-encoded IS1 *in vivo* (see Methods *Analysis of within-patient transposition events* for details) and for closing the reference genomes used in the RNA-Seq analysis.

### Genomic analyses

#### Quality control of Illumina reads

First, we carried out the quality control (QC) of the raw Illumina short reads using FastQC v.0.11.9, and merged all the reports with MultiQC v.1.11. We performed the trimming of the reads using Trim Galore v.0.6.4 or v0.6.6. We adjusted parameters to trim low-quality ends, discard short reads shorter than 50 bp and to trim possible adapters (-q 20 –length 50 –nextera). To check the trimming step and the quality of the trimmed reads, we performed a second QC over the trimmed reads, and used these for the following downstream analyses.

#### Reference genomes

To close the reference genomes of the ancestral strains (day 0) employed in the experimental evolution, the RNA-Seq and the within-patient analyses, we sequenced the stocked isolated strains carrying pOXA-48 both by short and long read sequencing as explained before. We performed hybrid assemblies of the short and long reads using Unicycler v0.4.9 or v.0.5.0 with default parameters and annotated the closed genomes using the NCBI Prokaryotic Genome Annotation Pipeline (PGAP; v2021-07-01.build5508 or v.2022-12-13.build6494) run locally. Strains which could not be closed by Unicycler were closed with Flye v2.9-b1768 and polished with Medaka v1.4.3 and several rounds of Pilon v1.24, mapping the short-reads (see Suppl. Table 1 and Code availability).

#### Analysis of mutations in the evolved populations

For the variant calling of short read data of the experimental ancestors (day 1 to avoid false positives) and the evolved populations (day 15), we used breseq v.0.36^45^. For population WGS data, we allowed the detection of polymorphic mutations (flag -p), and used the .gbk files generated by PGAP as reference for each strain. We analyzed the percentage of mapped reads of the evolved strains against the ancestral strains. All populations showed high levels of mapped reads indicating non-cross contaminations events during the EE (Suppl. Table 2). We discarded from further analysis a replicate of C309 bearing pOXA-48 in presence of AMC which showed 91% of mapped reads (Suppl. Fig. 7). To analyze the variant calling, we developed a Python script (Python v.3.8) to parse the HTML results given by breseq to obtain XLSX files (see Code availability section) and simultaneously compare mutations present in the samples by merging their reports into an individual table per strain. Essentially, this script takes the HTML tables reported by breseq (1 per replicate), and transforms their information into a spreadsheet table, considering the same mutations in multiple replicates, and merging them into the same entry. This is especially useful when analyzing numerous samples. However, the analysis of whole populations of clinical strains entails an additional challenge, as it increases the chances of reporting false positive mutations, specially when detecting polymorphisms^45^. To avoid this, we performed an exhaustive filtering step of the resulting mutations applying multiple conditions. Briefly, we: i) filtered out mutations present in the ancestral strains to avoid false positives occurring during the first overnight culture (day 1); ii) discarded mutations in genes with more than two events in the same replicate, as this is usually an indicative of a gene encoded in multiple copies in the genome; iii) established a frequency threshold of minimum 15% for both SNPs and NJs unless parallel evolution was detected; iv) filtered out New Junction events present in ancestors and in surrounding regions close to intergenic mutations (+/- 10 bp); and v) also considered that two NJ could belong to the same event (e.g. the 2 sides of an insertion) and merged them. As a control for the NJ detection, and following the recommendations of the breseq developers, we also ran breseq to predict mobile element insertions (annotating IS elements as repeat_region) using CF12 and K163 populations, for which we obtained the same results as in the previous analysis. To assess the mutation spectrum of each species, we summarized the SNPs and NJ events occurring in all of the strains and classified them depending on their class. SNPs were divided into intergenic, synonymous and nonsynonymous. We filtered out pseudogene mutations. For the visualization of the mutations in the genome of each strain replicates, we used the R package circlize v.0.4.15 (see Code Availability). Further, we identified IS-mediated NJ if one of the NJ extremes called an IS element, and classified them depending on the IS family.

#### Calculation of genomic IS1 copy number

We calculated the initial number of IS1 copies of the EE ancestors by summing the copies of annotated IS1 in the chromosome and in plasmids by PGAP. For plasmids, we corrected the number of annotated IS1 copies by the estimated copy number of the corresponding plasmid. We did not include the copies encoded in pOXA-48 in in the analysis of the total number of IS1 in the EE ancestors. We calculated the plasmid copy number (PCN) of all the plasmids from the mapping results as the ratio of each plasmid median sequencing coverage divided by the chromosome median coverage. We calculated the sequencing coverage at each position of the genome using Samtools v.1.13 (samtools depth). We estimated the median sequencing depth values using datamash v.1.4. For each replicate, we calculated the median sequencing depth for the chromosome, which shows one copy per cell. Then, to calculate the IS1 copy number, we corrected the number of copies encoded in plasmids by the respective plasmid copy number. For this, we divided the median sequencing depth of each plasmid and divided it by the median sequencing depth of the chromosome, thus obtaining a proxy of each plasmid copy number in the genome. A ratio of 1 indicates that chromosome and plasmid were sequenced at the same depth, and thus plasmid is present at roughly one copy per cell in the sample. Ratios above and below 1 would indicate increased copy number or loss of the plasmid in the sample, respectively.

#### Tracking IS1 elements

As the genomes from the different strains contained multiple copies of IS1 elements, we needed to confirm that the IS1 element was moving from pOXA-48 to the chromosome as breseq predicted. Moreover, the study of structural variants such as insertions caused by IS transpositions is intricate to assess with short-read data. Thus, we verified whether there were other identical copies of the IS1 element from pOXA-48 in the genome of each strain. Briefly, we extracted the sequence of the IS1 element from pOXA-48 (coord. *ca.* 30Kbp, identical to the one in the coord. *ca.* 10Kbp) using samtools faidx command (see Code Availability). Then, using BLASTn we looked for hits in the genome of each strain. We considered those hits which matched the length of the IS with 100% of identities to be the same IS1 element. Hence, if the reference genome contained only the copy of this specific IS1 in the pOXA-48 sequence, we would be able to track it down during the evolution, supporting the transposition from pOXA-48 to the chromosome.

#### Confirmation of structural variants

As with Illumina short-read data is difficult to determine large scale structural variants, we sequenced a subset of the evolved genomes were breseq results indicated pOXA-48-encoded IS1 insertions in the chromosome using Oxford Nanopore sequencing technologies (see WGS section; Suppl. Fig. 21). Specifically, we sequenced by both short and long reads 1-3 clones per evolved population of *K. pneumoniae* and *E. coli* carrying pOXA-48. We analyzed the short-read Illumina data as commented above, using breseq for the variant calling to confirm the presence of NJ of interest in the clones. Here, as we analyzed clonal samples, we run breseq with default parameters allowing for the detection of clonal mutations. For the analysis of long-read data, we performed the basecalling and demultiplexing of the raw ONT data using Guppy v.6.1.7+21b93d1a5. We used MinionQC to check the quality and length of the reads. Finally, we used Sniffles v.2.0.6 to call complex structural variants in the isolated clones. We combined these results with the breseq output both from the clonal and the population samples to confirm the insertions called by breseq.

#### Assembly of evolved clones

To confirm the insertion of IS1 during evolution, in addition to the variant calling with short and long read data, we performed hybrid assembly of the long and short read sequences for each clone per population evolved using Unicycler v.0.5.0 with default parameters. Then, we aligned the pOXA-48 IS1 sequence, extracted with samtools faidx command, using BLASTn against the closed genome. There, we confirmed again the NJ called by breseq in the polyclonal populations regarding pOXA-48 IS1 transposition.

#### Checking pOXA-48 sequence

To check that all the samples included in our study contained the most common variant of the pOXA-48 plasmid (K8), we checked the sequence of pOXA-48 in each of the ancestral strains. Using the short-read Illumina data of the ancestors, we performed the variant calling against the reference sequence of pOXA-48_K8^55^ using breseq v.0.36 with default parameters and allowing for polyclonal detection (flag -p). We considered all mutations which had a frequency of 100% (Suppl. Fig 21).

### Transcriptomic analyses

#### RNA extraction, sequencing and read quality control

We grew the cells in LB medium with continuous shaking. When cultures reached a turbidity of 0.5 at 600 nm, we collected the cells at 4°C, pelleted them by centrifugation at 4°C 12,000 g for 1 min and immediately froze them at -70°C. We purified the total RNA from each sample using the NZY Total RNA Isolation kit (NZYTECH). Then, we determined RNAs concentration using the Qubit RNA Broad-Range Assay following the manufacturer’s instructions. Additionally, the quality of the RNA was examined using the Tape-Station system (Agilent). We ribodepleted the RNA and sequenced it at the Wellcome Trust Centre for Human Genetics (WTCHG; Oxford, UK) using the Illumina’s NovaSeq6000 platform, resulting in >8 million reads per sample. We sequenced three biological replicates per strain and condition (carrying and not carrying pOXA-48), except for strains CF13, K209p and K209 that we sequenced two replicates due insufficient RNA Integrity number (RIN) (Suppl. Table 1). We trimmed and adapter-removed the reads with Trim Galore v0.6.4 (-q 20 --length 50 --illumina). We assessed the quality of the reads before and after trimming with FastQC v.0.11.9 and MultiQC v1.11.

#### RNA-Seq data analysis

We performed differential expression analysis to analyze the transcription of IS1 in a subset of strains carrying and not carrying pOXA-48. ISs typically bear multiple copies with high identity nucleotide sequences between them, and thus, short Illumina reads can map ambiguously to different IS copies in different locations of the genome, reducing the statistical power to detect differential expression of ISs between conditions. To avoid this, we first identified in the genomes all copies of all IS families annotated by PGAP. Then, we masked the sequences of all but one copy of ISs per family (leaving the unmasked single copy preferentially in the chromosome or in the largest plasmid) from the reference genomes, using the bedtools maskfasta tool v2.27.1. This way, reads belonging to an IS family (like IS1) can map with low ambiguity to one region of the genome. After re-annotating the masked genomes with PGAP v2021-07-01.build5508, we mapped the trimmed reads to their corresponding reference genome with BWA-MEM v0.7.17. The percentage of mapped reads was >99% for all samples. We inspected the mappings to confirm the appropriate presence or absence of pOXA-48 in the replicates. A replicate in K147c1 and in CF13c1 (cured from pOXA-48) actually carried the plasmid, so they were included as an additional replicate of the pOXA-48-carrying strains. We obtained the raw counts of reads mapping to each genomic feature (including CDS, ncRNA, tRNA, tmRNA, antisense RNA, RNase P and SRP RNA) with featureCounts from the Rsubread v2.14.2 package. We performed the differential expression analysis from raw counts using DESeq2 v1.40.1 by comparing the pOXA-48-carrying strain against the pOXA-48-free strain.

### Construction of pIS1, an inducible IS1-expressing plasmid

To construct pIS1, we exchanged the *gfp* gene from pBGA^55^ with the IS1 coding region from the *K. pneumoniae* strain K163. pBGA is a ColE1-type plasmid with ∼19 copies per cell^9,55^. In pIS1, the IS1 gene is under the control of the P_BAD_ promoter, so the IS1 production is repressed and selectively induced by the presence of L-arabinose. Importantly, the two inverted repeat regions (IR) flanking the transposase gene are not present in pIS1. The excluded IR sequences are 5’-GGTAATGACTCCAACTTATTGAT-3’ and 5’-GGTGATGCTGCCAACTTACTGA-3’, which we identified using ISEScan v.1.7.2.3. We used primers ID-1 and ID-2 to amplify IS1 and ID-3 and ID-4 primers to amplify the pBGA backbone (Phusion Hot Start II DNA Polymerase, ThermoFisher Scientific, MA, USA). All primer sequences used in this study are listed in Suppl. Table 7. Then, we purified the PCR products (NucleoSpin PCR clean-up, Macherey-Nagel, USA) and digested them with DpnI (New England BioLabs, MA, USA). We constructed pIS1 by joining the fragments using the Gibson Assembly Cloning kit (New England BioLabs, MA, USA). We transformed the resulting reaction into *E. coli* NEB 10-beta (New England BioLabs, MA, USA) by electroporation, following the manufacturer’s recommendations. Later, we selected pIS1 transformants on LB agar plates supplemented with apramycin (30 mg/mL) and verified transformants by PCR using primers ID-5 and ID-6. Then, we extracted pIS1 plasmids (NucleoSpin Plasmid EasyPure, Macherey-Nagel, USA) from bacterial cells and validated the pIS1 sequence by sequencing (Oxford Nanopore in Plasmidsaurus, OR, USA). Finally, we inoculated 500 mL Erlenmeyer flasks filled with 100 mL of LB with overnight cultures of pOXA-48-carrying and pOXA-48-free K25. We incubated them at 37°C with continuous agitation (250 r.p.m., Thermo Scientific™ MaxQ™ 8000). After two hours of incubation, we transferred the cultures to 50 mL tubes and centrifuged them at 4°C (3,000 g, 15 minutes). We discarded the supernatant and washed the pellets three times with cold sterile water (Thermo Fisher Scientific, MA, USA). After that, we transformed pIS1 into each phenotype by electroporation using 0.5 cm cuvettes and a 2.5 kV pulse (MicroPulser Electroporator, Biorad Spain). We selected transformants on LB agar plates supplemented with apramycin 30 mg/mL and we verified the presence of pIS1 by PCR as previously described. We stored overnight cultures of each genotype in LB (15% glycerol) in cryotubes at -80°C.

### Fluctuation assay to test IS1 transposition rate

We conducted fluctuation assays using the following strains: K25, K25 with pOXA-48, K25 with pIS1 and K25 with pOXA-48 and pIS1. We inoculated independent cultures of each strain allowing them to grow at 37°C for 22 h with a constant shaking of 225 r.p.m. To estimate if the presence of pOXA-48 affected the mutation rate we inoculated 80 independent cultures of K25 and K25 with pOXA-48 in LB. To test if the InsA protein repressed the transposition of the pOXA-48 encoded plasmid, we inoculated 20 independent cultures of K25 with pOXA-48 and pIS1 in LB with 0.5% of L-arabinose and 20 without L-arabinose. The presence of L-arabinose induced the expression of the pIS1-encoded *insAB* operon. To discard any effect of the L-arabinose in the loss-of-capsule mutation rate, we also inoculated 20 independent cultures of K25, K25 with pIS1 and K25 with pOXA-48 with and without 0.5% of L-arabinose. Subsequently, we plated aliquotes in LB-Agar plates to quantify the spontaneous mutants with loss-of-capsule phenotype (Fig. 4C). We counted the colonies and determined the total population size and the fraction of capsule-free mutants based on phenotypic differences (Fig. 4C). Using the rSalvador package, we estimated the phenotypic mutation rate for the loss-of-capsule phenotype^71^. To analyze the genetic changes responsible for this phenotype, we conducted whole-genome sequencing using the NextSeq 2000 platform at SeqCoast Genomics (Portsmouth, NH, USA) on 32 capsule-free strains: 17 strains of K25 with pOXA, 12 strains of K25, 5 strains of K25 with pOXA-48 and pIS1 grown in the absence of arabinose and 5 strains of K25 with pOXA-48 and pIS1 grown in the presence of arabinose.

### Relative fitness determination by competition assays

To test the fitness effects associated with the loss of capsule, we selected 6 capsule-free independent clones from the fluctuation assays. We selected three capsule-free colonies from the K25 genotype (clones 81, 82 and 92) and three from the K25 with pOXA-48 genotype (clones 105, 107 and 110) (Suppl. Fig. 18C and Suppl. Table 5). We then competed the clones 81, 82 and 92 against K25, and the clones 105, 107 and 110 against K25 with pOXA-48. We selected each competitor from individual colonies isolated in LB agar plates, and resuspended in NaCl 0.9% solution to an OD 600nm of 1. Then, we mixed each competitor (1:1) and we diluted 400-fold the mixture in 200µl of fresh LB in 96-well plates and incubated for 20 h at 37°C and 365 r.p.m. We plated in LB-Agar plates appropriate dilutions of each competition immediately after mixing and after 20 hours of growth and we estimated the number of colonies expressing and lacking the capsule. We calculated the relative fitness using the formula:

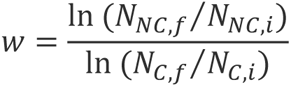

where *w* is the relative fitness of the capsule-free clones compared to the capsulated wild-type, *N_C_* (capsulated) and *N_NC_* (Non-capsulated) are the number of cells expressing or not the capsule respectively. *N_i_* and *N_f_* indicates the number of cells at the beginning and at the end of the competition respectively. Each competition was done by triplicate.

### Analysis of within-patient transposition events

#### Variant calling of in vivo evolved clones

To analyze whether our results could be translated to real-life scenarios, we decided to study the possibility of transposition events of IS1 occurring in the presence of pOXA-48 in the gut of hospitalized patients. For this, we selected strains from the R-GNOSIS collection with multiple isolates from the same patient. Thus, we expected to observe *in vivo* events of evolution. Specifically, we analyzed 72 isolates (21 *E. coli*, 2 *C. freundii*, 49 *K. pneumoniae*), from 31 strains (10 *E. coli*, 1 *C. freundii*, 20 *K. pneumoniae*) and 27 different patients. We performed the hybrid assembly of one isolate for each of the strains using Unicycler or Flye+Medaka+Pilon as previously explained, and annotated the assemblies with PGAP. We later used these assemblies as reference to analyze the mutations in the respective evolved clones using breseq. We used the same filtering criteria as for the EE genomic analyses to clean the resulting mutations. 2 of the isolates did not fulfill our criteria for being considered the same clones, as they showed >7 SNPs, and we kept them out of the analyses.

#### General Additive Model (GAM) of IS1 transposition in vivo

We built a generalized additive model (GAM) to study the effects of genomic IS1 copy number and the bacterial species on IS1 transposition rate, using the mgcv v.1.8-42 package in R. We calculated the IS1 transposition rate *in vivo* as the ratio between the number of IS1 transposition events that occured between two subsequent isolations of the same bacterial lineage, and the days spanned between the isolations. We included the genomic IS1 copy number of each strain as a smooth term, and the species factor as a categorical term (See Code Availability, GAM model function: gam(IS1_per_day ∼ s(nIS1_genome, k = 6) + Species, data = summary_patients_table, method = “REML”)). Additionally, we controlled for the complexity of the model varying the k parameter (number of basis functions) and selected the optimal one (k = 6).

### Statistical analysis

We performed all the statistical analyses in Rstudio v.2022.02.3+492 (R v.4.2.2 patched 2022-11-10 r83330). We tested the normality and homoscedasticity of the data using Shapiro-Wilk and Bartlett or Levene tests. Depending on the structure of the data, we applied parametric or non-parametric tests for the comparison of multiple groups. For normal data, we applied ANOVA tests, and for non-normal data, Kruskal-Wallis tests. For multiple comparisons between more than 2 groups, we used Tukey tests, after ANOVA, or Mann-Whitney tests, after Kruskal-Wallis, as post-hoc analysis. We used Pearson’s correlation when data was normal, and Spearman’s with non-normal data.

## Data availability

The sequences generated for the EE, analyzed during this work, can be found at the Sequence Read Archive (SRA) repository under the BioProject ID: <u>PRJNA1076826</u>. The genomic sequences used for the study of the within-patient evolution section in this project were previously generated and uploaded at the SRA repository under the BioProject ID: <u>PRJNA626430</u>. Reference genomes that were previously generated are available under the BioProjects <u>PRJNA626430</u> and <u>PRJNA838107</u>. For closing certain reference genomes, Illumina reads from BioProject <u>PRJNA641166</u> were used (Suppl. Table 1).

## Code availability

All the code developed for the analyses included in this work can be found at the Github repository https://github.com/jorgEVOplasmids/rapid_adaptation_pOXA48. The extended code for the RNA-Seq analysis can be found at the GitHub repository https://github.com/LaboraTORIbio/RNA-Seq_enterobacteria_pOXA-48.

## Acknowledgements

We thank J.A. Escudero for constructive comments on the manuscript, O. Rendueles for helping discussion regarding *K. pneumoniae* capsules, C. W. Marshall for bioinformatic support, F. Trigo da Roza for helping with Long-Read sequencing and L. Jaraba and A. Alonso-del Valle for technical support. This work was supported by the European Research Council (ERC) under the European Union’s Horizon 2020 research and innovation programme (ERC grant no. 757440-PLASREVOLUTION, to A.S.M), by the Instituto de Salud Carlos III (PI19/00749, to A.S.M) cofunded by the European Development Regional Fund ‘A way to achieve Europe’, by the “La Caixa’’ Foundation (ID 100010434) under the project (LCF/BQ/PR22/11920001, to A.S.L), by the European Commission (nos. H2020-MSCA-IF-2019, 895671-REPLAY, to A.S.L) and by the European Society of Clinical Microbiology and Infectious Diseases (Research Grant 2022, to A.S.L.). JR-B acknowledges support by a Miguel Servet contract from the Carlos III Health Institute (ISCIII) (grant no. CP20/00154), co-founded by the European Social Fund, ‘Investing in your future’, PI21/01363, funded by the Carlos III Health Institute (ISCIII) and co-funded by the European Union and the European Union (ERC, HorizonGT, 101077809). R.C. research is supported by CIBER de Enfermedades Infecciosas (CIBERINFEC) (CB21/13/00081, CB21/13/00084), Instituto de Salud Carlos III, Madrid, Spain. M.H.G. is supported by a postdoctoral contract by CIBERINFEC (CB21/13/00084). C.H. is supported by a Sara Borrel contract from the Instituto de Salud Carlos III (ISCIII) (grant no. CD21/00115), the Convocatoria Intramural Emergentes 2021 FIBioHRC-IRYCIS. Cod. IMP-21 n° C13, and PI23/01945 project funded by Carlos III Health Institute (ISCIII).

## Author contributions

J.S.D., A.S.L. and A.S.M conceptualized the study. J.S.D., A.S.L. and A.S.M. designed the methodology. J.S.D. analyzed the genomic data. A.S.L., J.D.F., C.H., C.C., P.H.G. and J.R.B. performed the experiments. J.D.F. and L.T.C. contributed to data analysis. M.H.G and R.C. designed and supervised sampling and the collection of bacterial isolates. J.S.D., A.S.L and A.S.M. analyzed the data and prepared the original draft of the manuscript and undertook the reviewing and editing. A.S.L. and A.S.M. were responsible for funding acquisition and supervision.

## Supplementary material

**Supplementary Figure 1.**
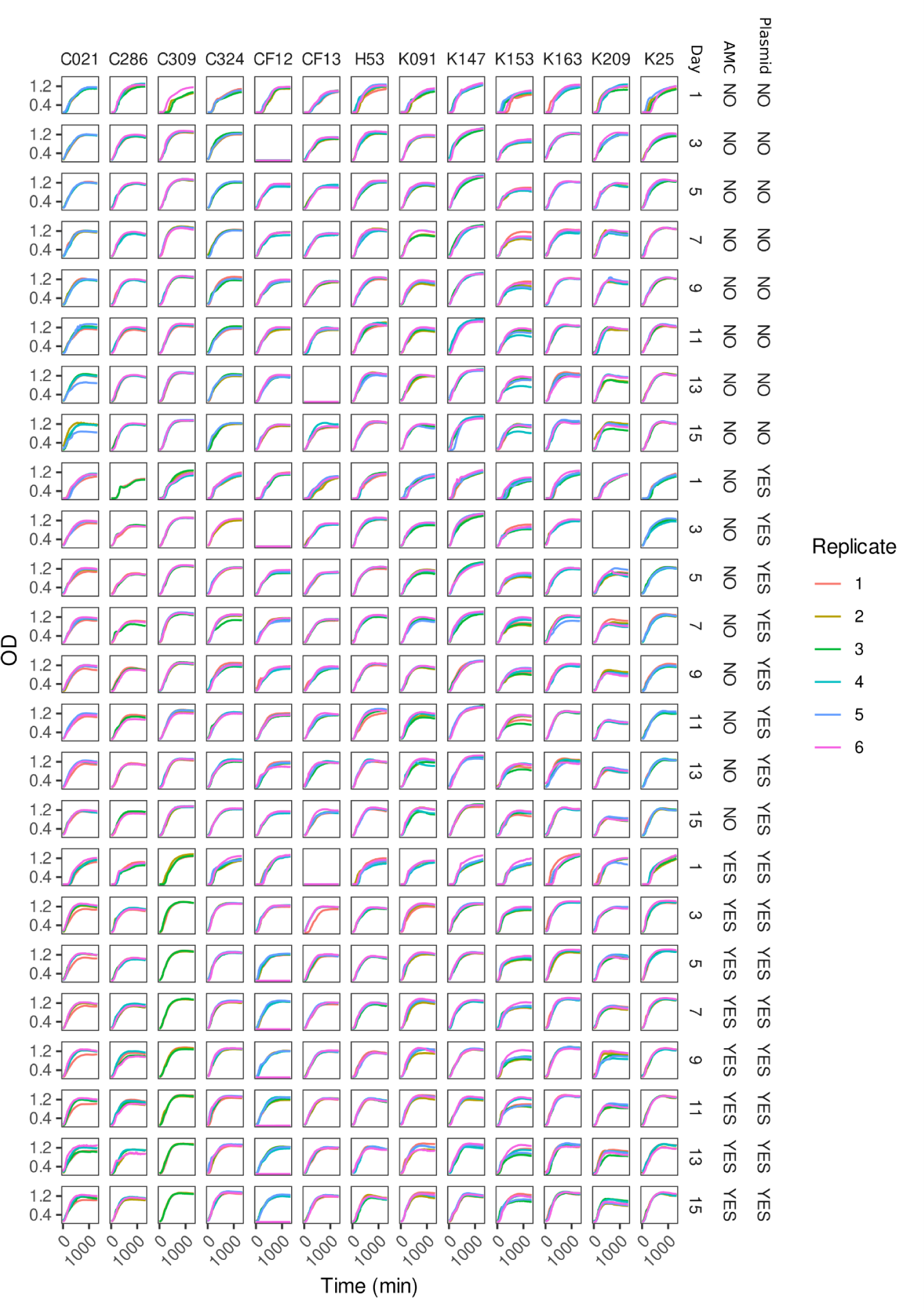
Growth curves during EE. Optical Density at 600 nm wavelength (OD600) of the strains included in this study measured during 24 hours in alternative days during the EE assay. The day of the EE assay in which we measured bacterial growth is indicated on the right, as well as the experimental conditions (presence/absence of pOXA-48 and presence/absence of AMC). Each strain name is indicated on the top. Each color of the curves represents a different replicate for each strain and condition.

**Supplementary Figure 2.**
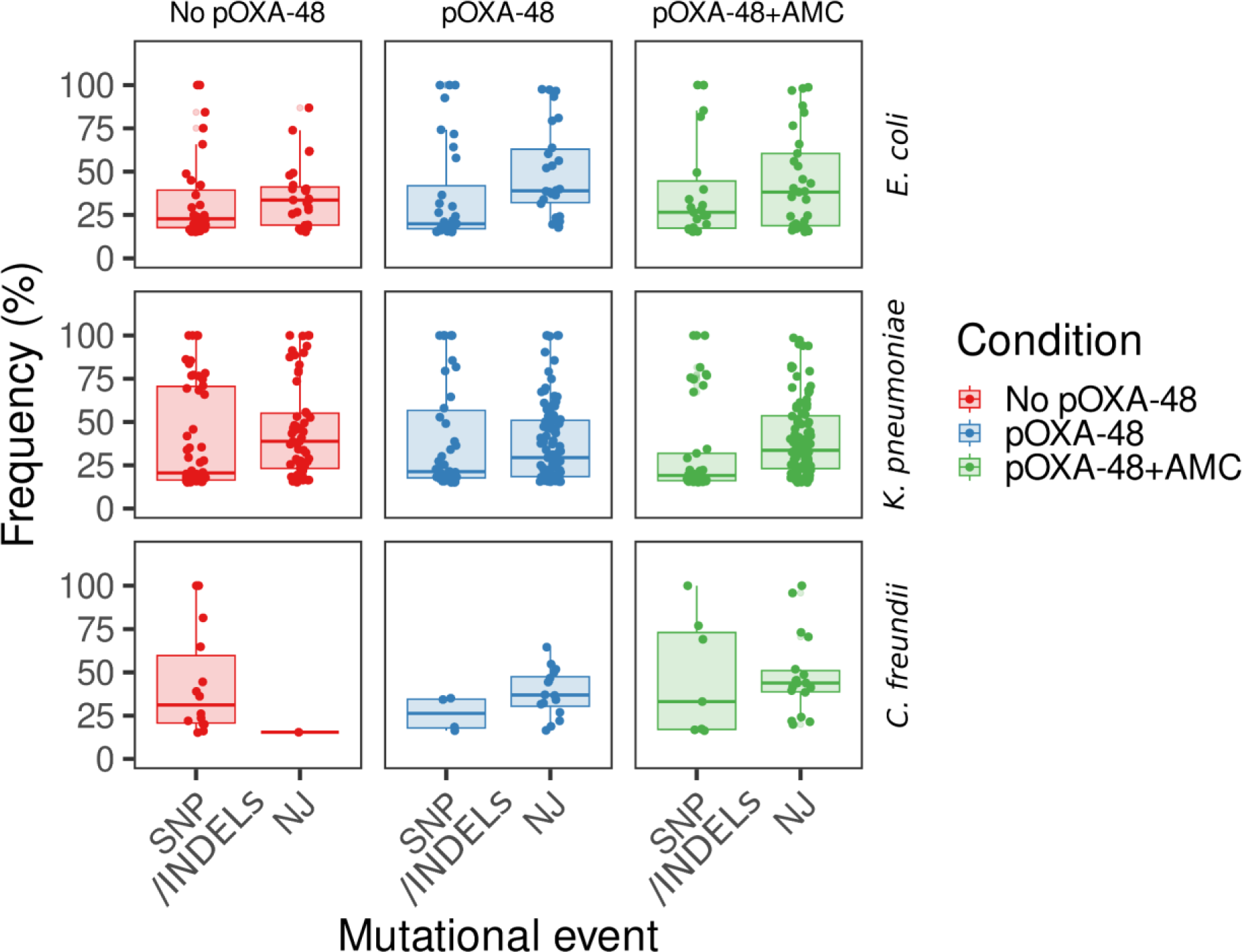
Frequency of New Junction events (NJ) and Single Nucleotide Polymorphisms (SNPs) or small insertions or deletions (INDELs) during the EE in the different conditions. NJ events are mutations predicted from split-read sequences which match distant locations in the reference sequence, such as large (>150 bp) deletions/insertions^45^. The frequency of each of the mutational events is shown for each species and experimental condition at the end of the EE. In general, NJs showed high frequencies for all the species and conditions, suggesting that these potentially produced more notable effects on bacterial fitness during the experiment. From all the mutations captured, very few were fixed (100% freq.; n=26/256 (10.2%) of the total SNPs; n=27/376 (7.1%) of the total NJs) in the populations at the end of the experiment.

**Supplementary Figure 3.**
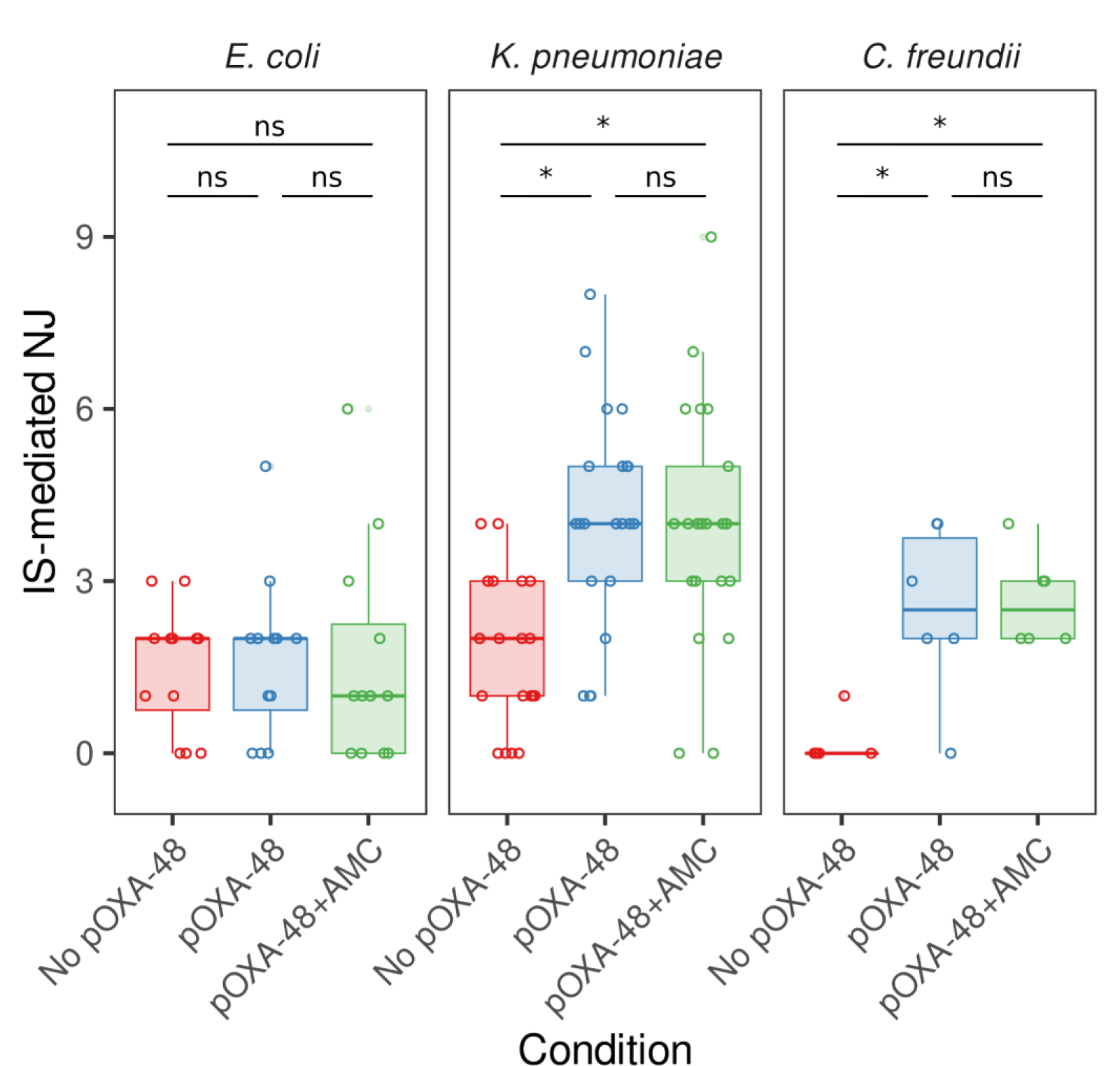
Number of IS-mediated NJ events during the EE in different conditions. The number of NJ events mediated by IS elements at day 15 of the EE for each species and experimental condition is shown. Each point represents an individual replicate population. Statistical comparison between the movement of ISs in the different conditions in each of the species revealed a significant increase in IS movements when the plasmid was present during the evolution in *K. pneumoniae* and *C. freundii* strains (two-sided Kruskal-Wallis test, p < 0.01) chi-squared = 18.551, d.f. = 2, p = 9.369e-05 for *K. pneumoniae*; chi-squared = 9.9479, d.f. = 2, p = 0.006916 for *C. freundii*, followed by two sided pairwise comparisons using Wilcoxon rank-sum test p > 0.05), but not in *E. coli* (two-sided Kruskal-Wallis test, chi-squared = 0.33163, d.f. = 2, p = 0.8472, followed by two-sided pairwise comparisons using Wilcoxon rank-sum test p > 0.05).

**Supplementary Figure 4.**
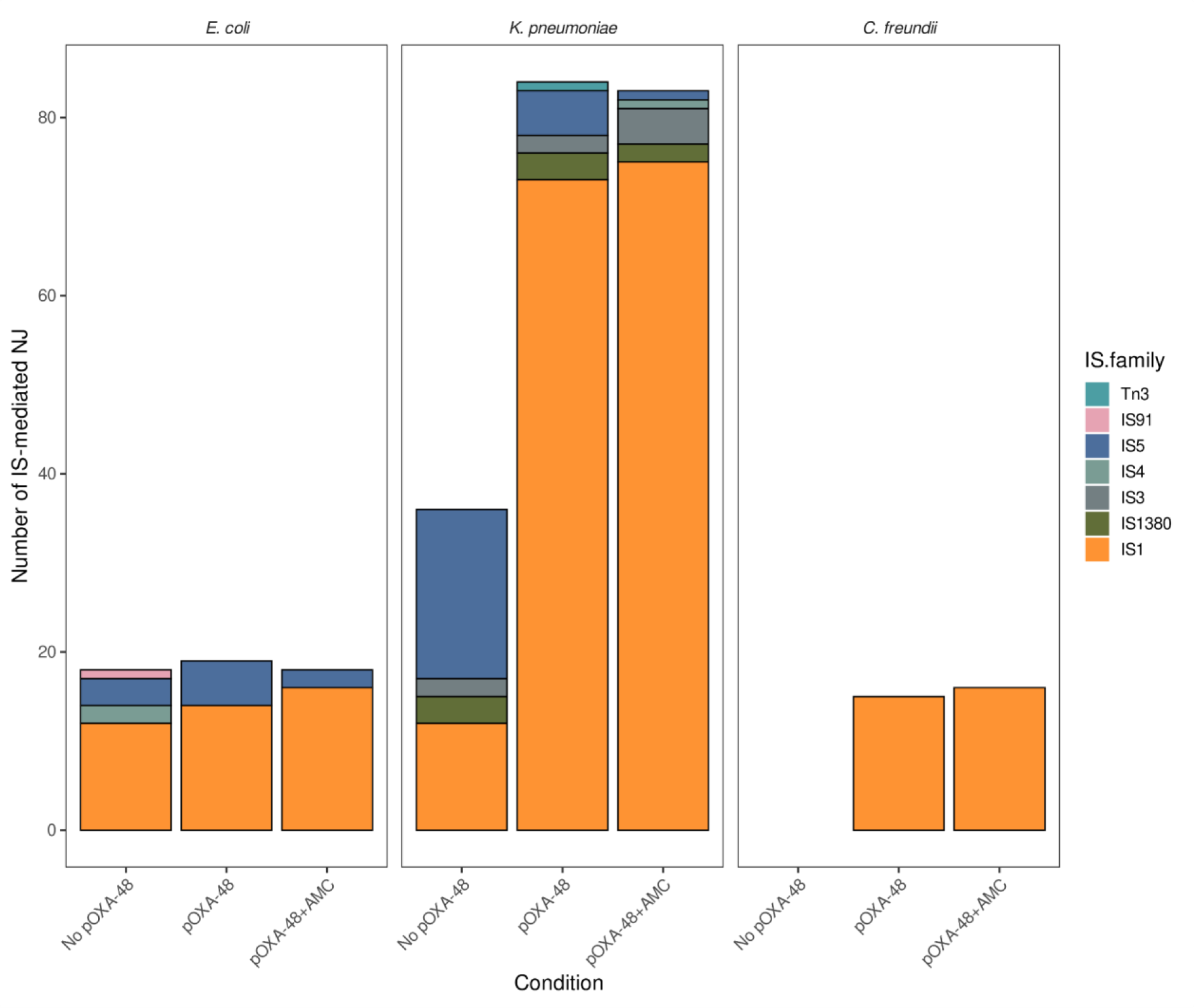
IS families transposition during EE. Number of transposition events associated with each of the IS families during the evolution for each species and experimental condition. Barplots represent the number of transposition events per IS family summarized for all the strains of each species studied. We identified those NJ events mediated by ISs in each population, and classified them depending on the family. We noticed that IS1 elements showed specially high transposition levels in most of the species and conditions. However, both in *C. freundii* and *K. pneumoniae* we observed that the presence of pOXA-48 significantly triggered the mobilization of IS1 during evolution (stats in Suppl. Fig. 3).

**Supplementary Figure 5.**
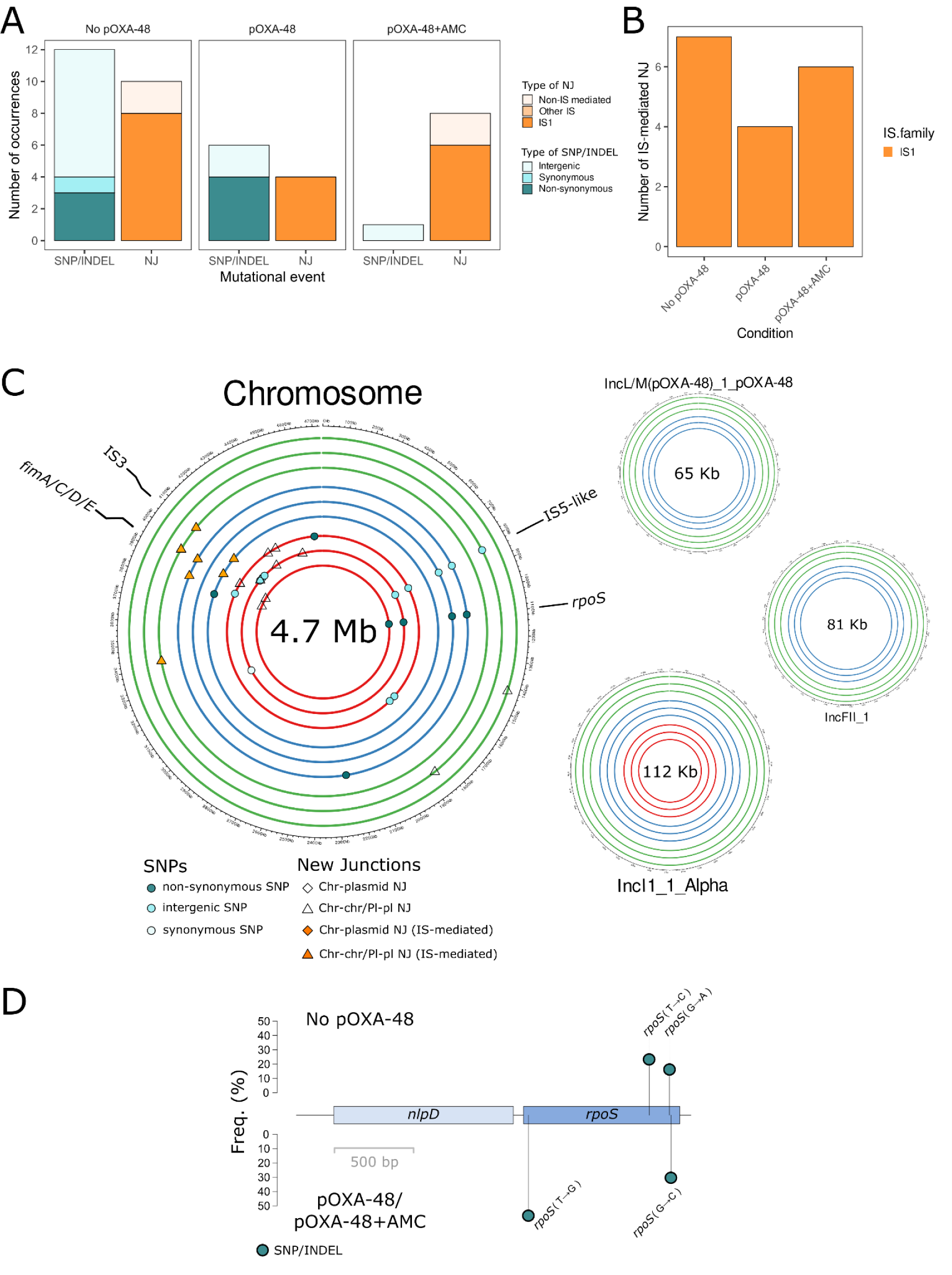
Summary of genomic changes during EE in strain C021 (*E. coli* ST10). **A)** Summary of the genetic changes in the propagated populations at day 15 classified in SNPs/INDELs or NJ for each experimental condition. Barplot colors indicate the type of event. **B)** NJ events classified by IS families per EE condition. **C)** Circa plot of the C021 genome showing the location of the genomic changes. From inside to outside, circles indicate the different replicates of the evolved bacterial populations without pOXA-48 (red), carrying pOXA-48 (blue) and carrying pOXA-48 in presence of AMC (green). Parallel evolution targets are labeled in each circa plot. The different types of SNPs/INDELs are represented by dots, whereas NJ events are depicted by shapes, filled in orange in the case of IS-mediated NJs. *rpoS* and the fimbrial operon were the main targets during the EE. All replicates without pOXA-48 also lost the IncFII plasmid. **D)** Lolliplot of the *rpoS-nlpD* stress response operon. The top part of the plot shows mutations occurred during the EE in the replicates propagated in absence of the plasmid, whereas the bottom part includes the events that happened in pOXA-48 carrying replicates both with and without AMC. Names of the genes are shown. The pathways of adaptation differed between pOXA-48-free and pOXA-48-carrying populations, with IS1 knocking out *rpoS* in pOXA-48-carrying populations.

**Supplementary Figure 6.**
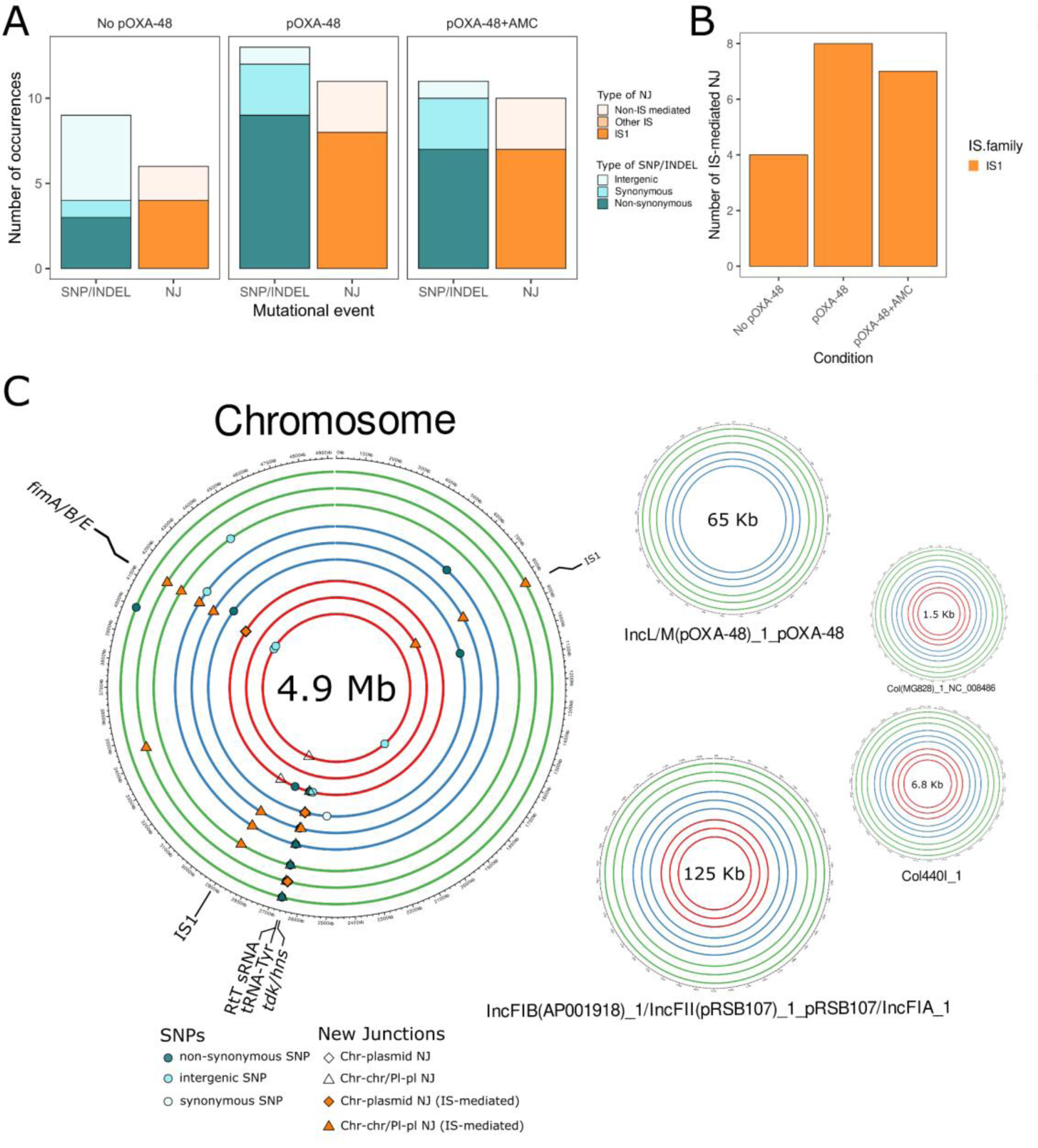
Summary of genomic changes during EE in strain C286 (*E. coli* ST405). **A)** Summary of the genetic changes in the propagated populations at day 15 classified in SNPs/INDELs or NJ for each experimental condition. Barplot colors indicate the type of event. **B)** NJ events classified by IS families per EE condition. **C)** Circa plots of the C286 genome showing the location of the genomic changes. From inside to outside, circles indicate the different replicates of the evolved bacterial populations without pOXA-48 (red), carrying pOXA-48 (blue) and carrying pOXA-48 in presence of AMC (green). Parallel evolution targets are labeled in each circa plot. The different types of SNPs/INDELs are represented by dots, whereas NJ events are depicted by shapes, filled in orange in the case of IS-mediated NJs. The fimbrial operon was one of the main targets during the EE.

**Supplementary Figure 7.**
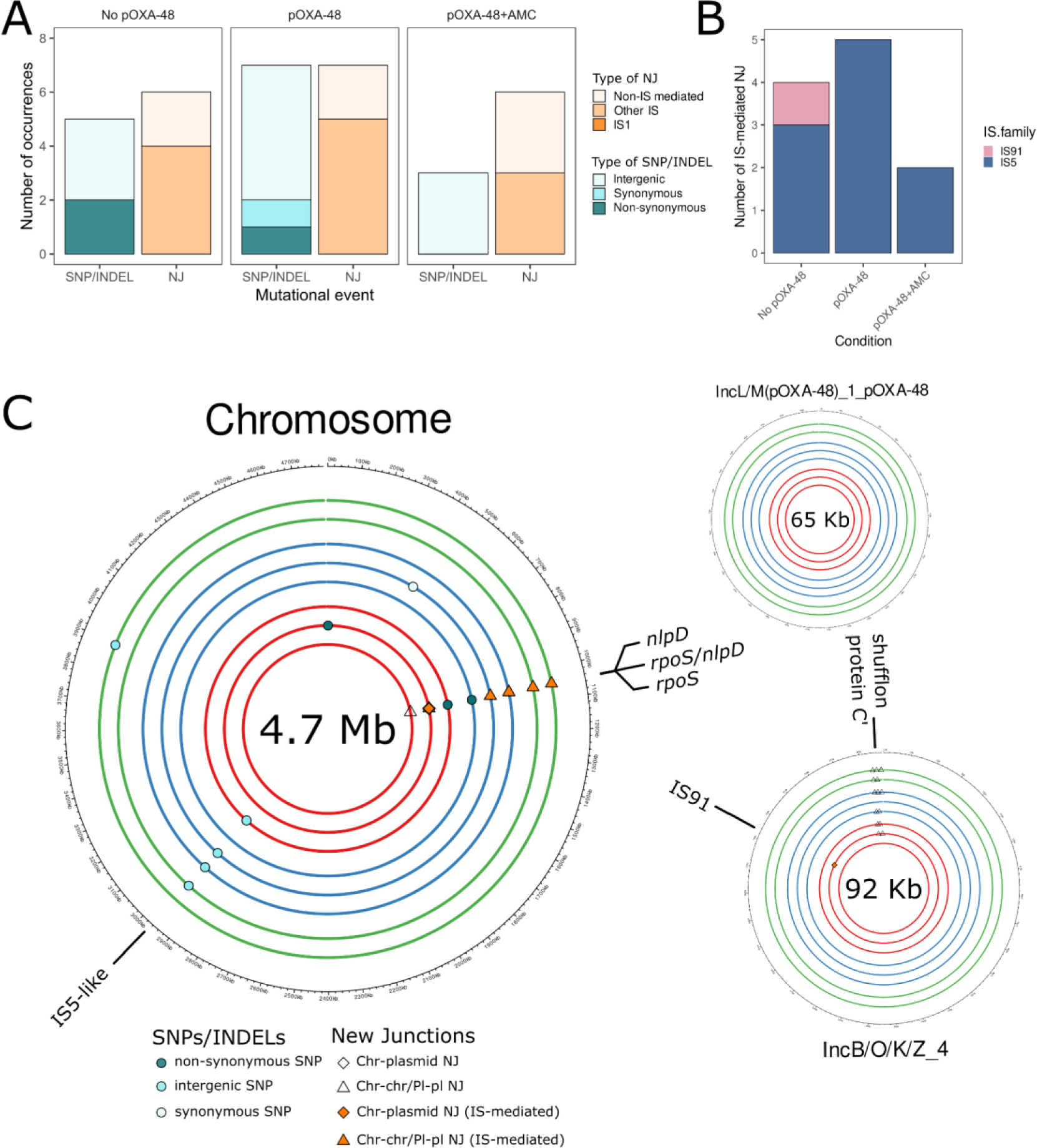
Summary of genomic changes during EE in strain C309 (*E. coli* ST6217). **A)** Summary of the genetic changes in the propagated populations at day 15 classified in SNPs/INDELs or NJ for each experimental condition. Barplot colors indicate the type of event. One of the 3 replicates with pOXA-48+AMC showed possible traces of cross-contamination, and therefore was eliminated from further analyses. **B)** NJ events classified by IS families per EE condition. In this strain we observed no transposition of IS1, but of IS5 and IS91. **C)** Circa plots of the C309 genome showing the location of the genomic changes. From inside to outside, circles indicate the different replicates of the evolved bacterial populations without pOXA-48 (red), carrying pOXA-48 (blue) and carrying pOXA-48 in presence of AMC (green). Parallel evolution targets are labeled in each circa plot. The different types of SNPs/INDELs are represented by dots, whereas NJ events are depicted by shapes, filled in orange in the case of IS-mediated NJs. *rpoS* and the fimbrial operon were the main targets during the EE. The *rpoS-nlpD* stress response operon was the main target of adaptation. In this case, the IS families commented in the B) panel mediated the knock-out of these genes both in presence and absence of pOXA-48, as well as some SNPs.

**Supplementary Figure 8.**
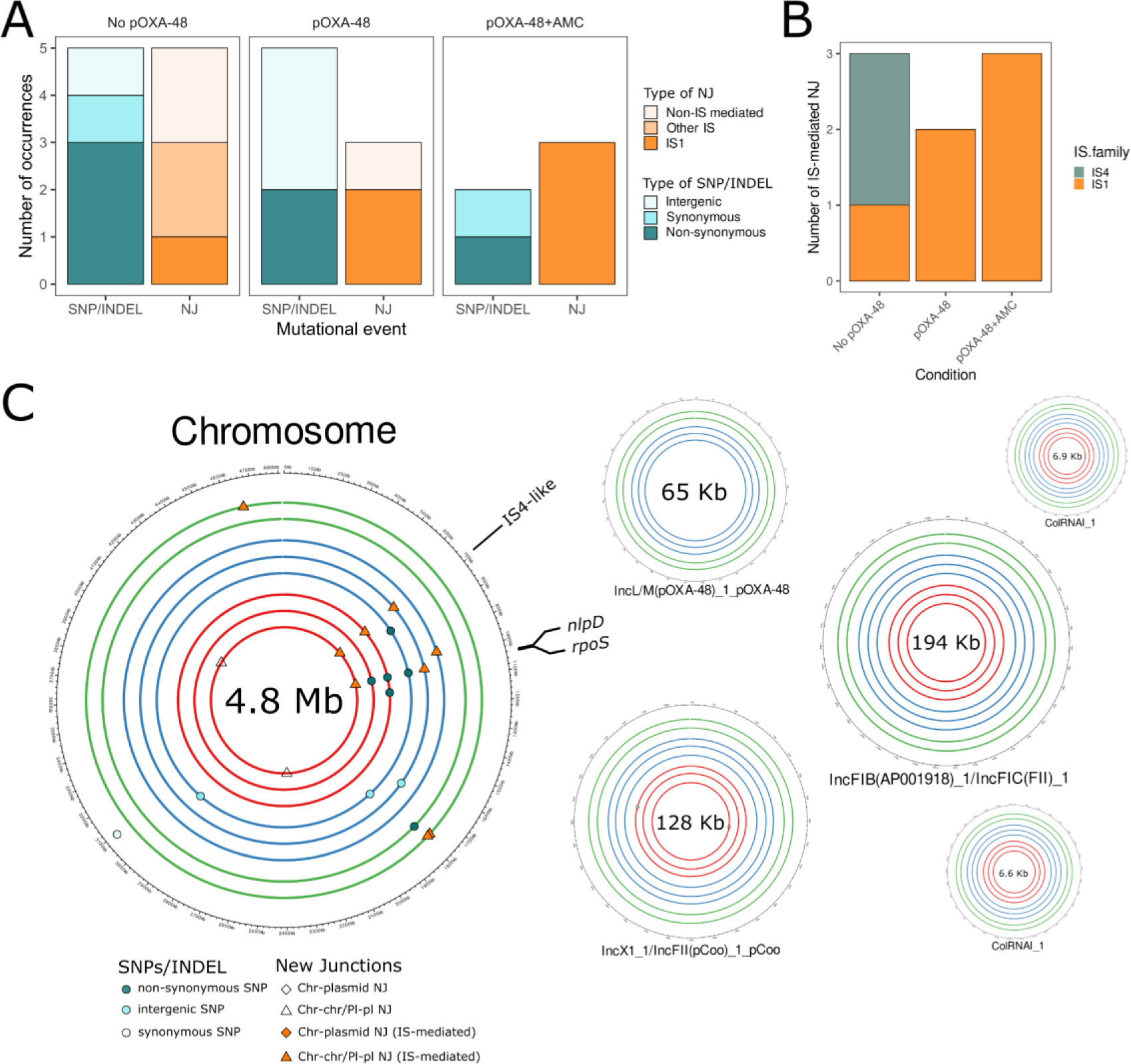
Summary of genomic changes during EE in strain C324 (*E. coli* ST10). **A)** Summary of the genetic changes in the propagated populations at day 15 classified in SNPs/INDELs or NJ for each experimental condition. Barplot colors indicate the type of event. **B)** NJ events classified by IS families per EE condition. In this case, IS4 and IS1 were the only families detected. **C)** Circa plots of the C324 genome showing the location of the genomic changes. From inside to outside, circles indicate the different replicates of the evolved bacterial populations without pOXA-48 (red), carrying pOXA-48 (blue) and carrying pOXA-48 in presence of AMC (green). Parallel evolution targets are labeled in each circa plot. The different types of SNPs/INDELs are represented by dots, whereas NJ events are depicted by shapes, filled in orange in the case of IS-mediated NJs. The *rpoS-nlpD* stress response operon was the main target of adaptation. We kept out of the analyses the hypermutator replicate (D5 mutated in *mutS*) propagated with pOXA-48 and AMC.

**Supplementary Figure 9.**
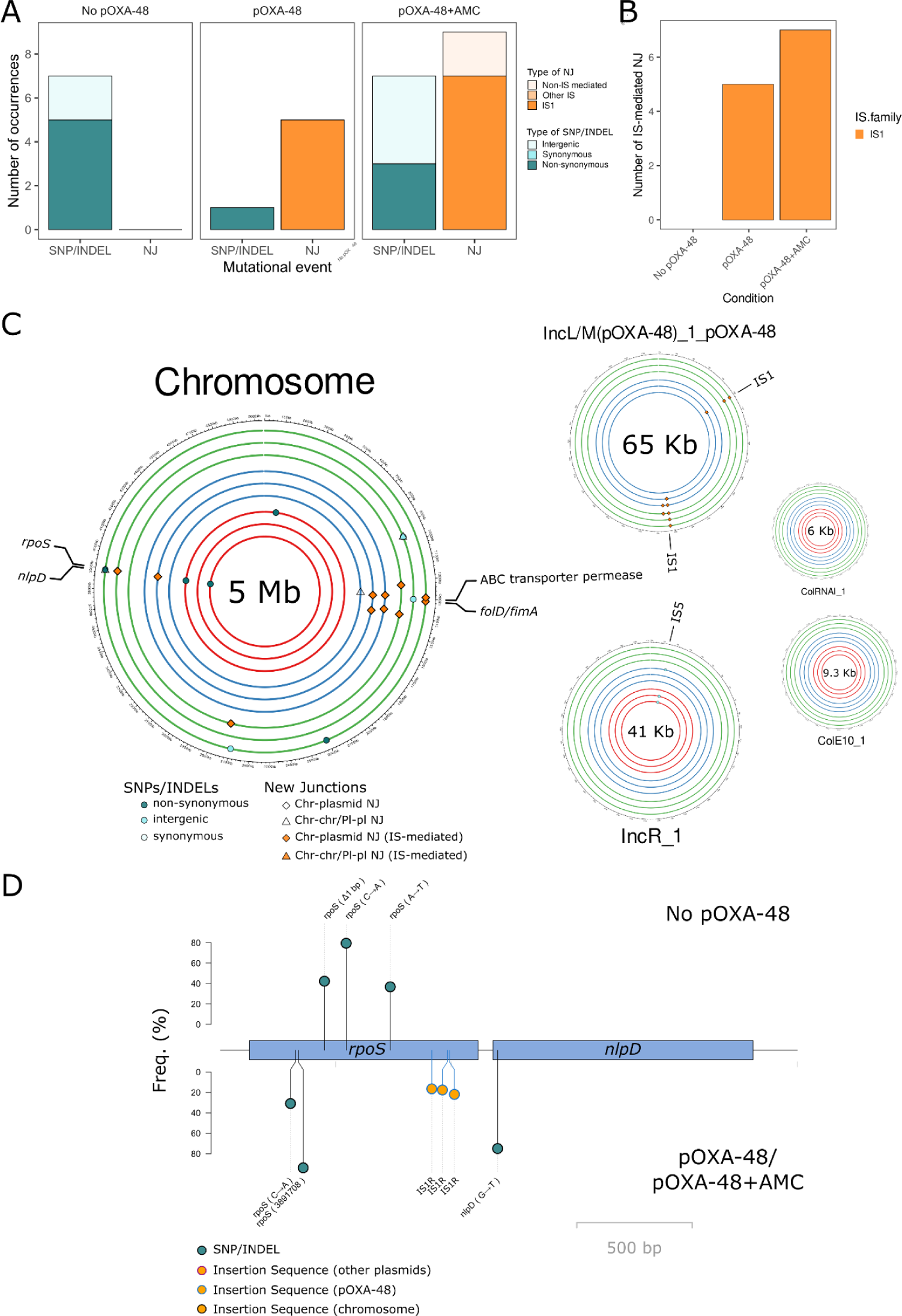
Summary of genomic changes during EE in strain CF12 (*C. freundii* ST22). **A)** Summary of the genetic changes in the propagated populations at day 15 classified in SNPs/INDELs or NJ for each experimental condition. Barplot colors indicate the type of event. **B)** NJ events classified by IS families per EE condition. IS1 transposition events were triggered by pOXA-48 presence during the EE. **C)** Circa plots of the CF12 genome showing the location of the genomic changes. From inside to outside, circles indicate the different replicates of the evolved bacterial populations without pOXA-48 (red), carrying pOXA-48 (blue) and carrying pOXA-48 in presence of AMC (green). Parallel evolution targets are labeled in each circa plot. The different types of SNPs/INDELs are represented by dots, whereas NJ events are depicted by shapes, filled in orange in the case of IS-mediated NJs. The *rpoS-nlpD* stress response operon and the fimbrial operon were the main target of adaptation. **D)** Lolliplot of the *rpoS-nlpD* stress response operon. The top part of the plot shows mutations occurred during the EE in the replicates propagated in absence of the plasmid, whereas the bottom part includes the events that happened in pOXA-48 carrying replicates both with and without AMC. Names of the genes are shown. The pathways of adaptation differed between pOXA-48-free and pOXA-48-carrying populations, with IS1 knocking out *rpoS* in pOXA-48-carrying populations.

**Supplementary Figure 10.**
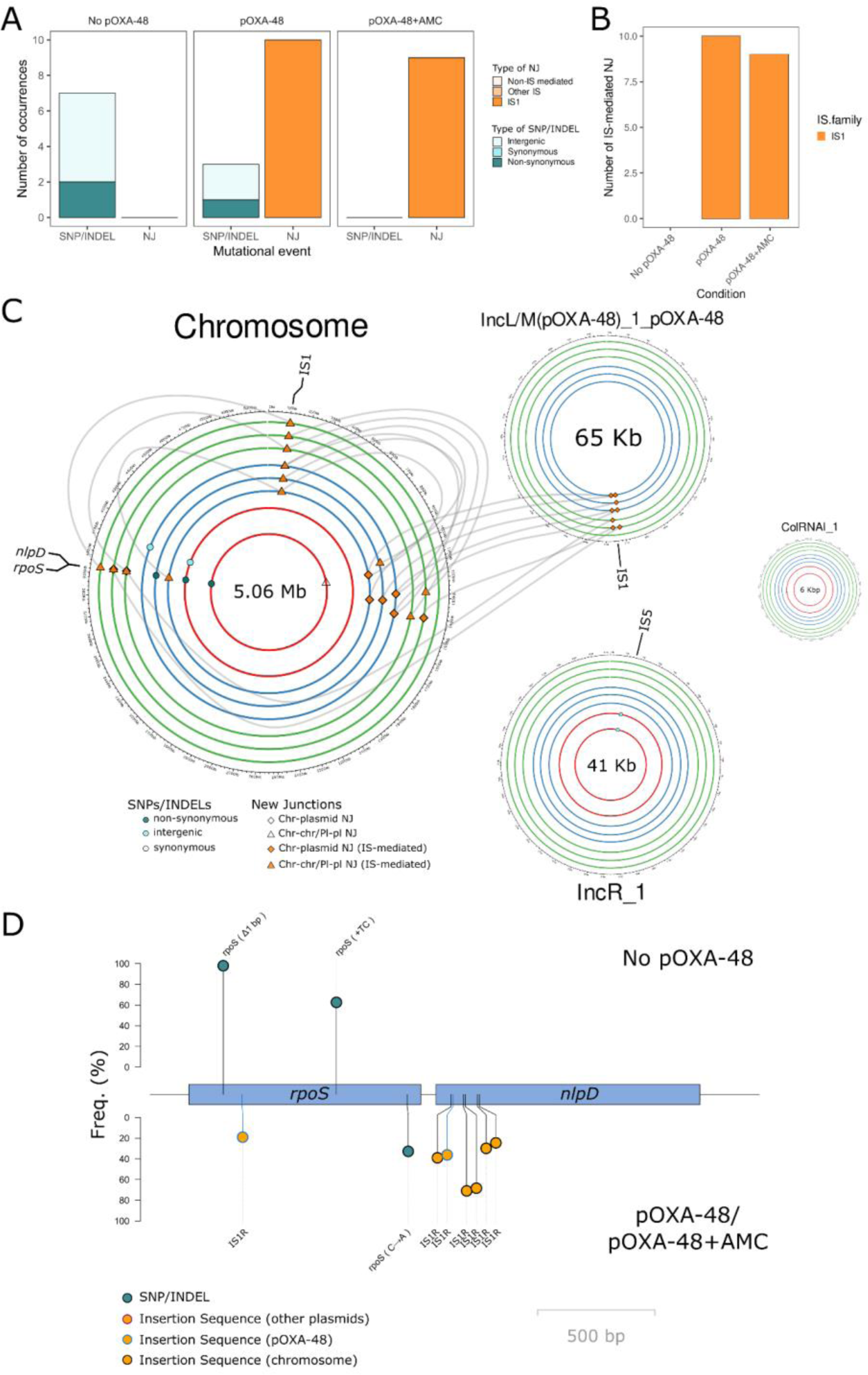
Summary of genomic changes during EE in strain CF13 (*C. freundii* ST22). **A)** Summary of the genetic changes in the propagated populations at day 15 classified in SNPs/INDELs or NJ for each experimental condition. Barplot colors indicate the type of event. **B)** NJ events classified by IS families per EE condition. IS1 transposition events were triggered by pOXA-48 presence during the EE. **C)** Circa plots of the CF13 genome showing the location of the genomic changes. From inside to outside, circles indicate the different replicates of the evolved bacterial populations without pOXA-48 (red), carrying pOXA-48 (blue) and carrying pOXA-48 in presence of AMC (green). Parallel evolution targets are labeled in each circa plot. The different types of SNPs/INDELs are represented by dots, whereas NJ events are depicted by shapes, filled in orange in the case of IS-mediated NJs. The *rpoS-nlpD* stress response operon and the fimbrial operon were the main target of adaptation. We kept out of the analyses a contaminated replicate propagated without pOXA-48. **D)** Lolliplot of the *rpoS-nlpD* stress response operon. The top part of the plot shows mutations occurred during the EE in the replicates propagated in absence of the plasmid, whereas the bottom part includes the events that happened in pOXA-48 carrying replicates both with and without AMC. Names of the genes are shown. The pathways of adaptation differed between pOXA-48-free and pOXA-48-carrying populations, with IS1 knocking out *rpoS* and *nlpD* in pOXA-48-carrying populations.

**Supplementary Figure 11.**
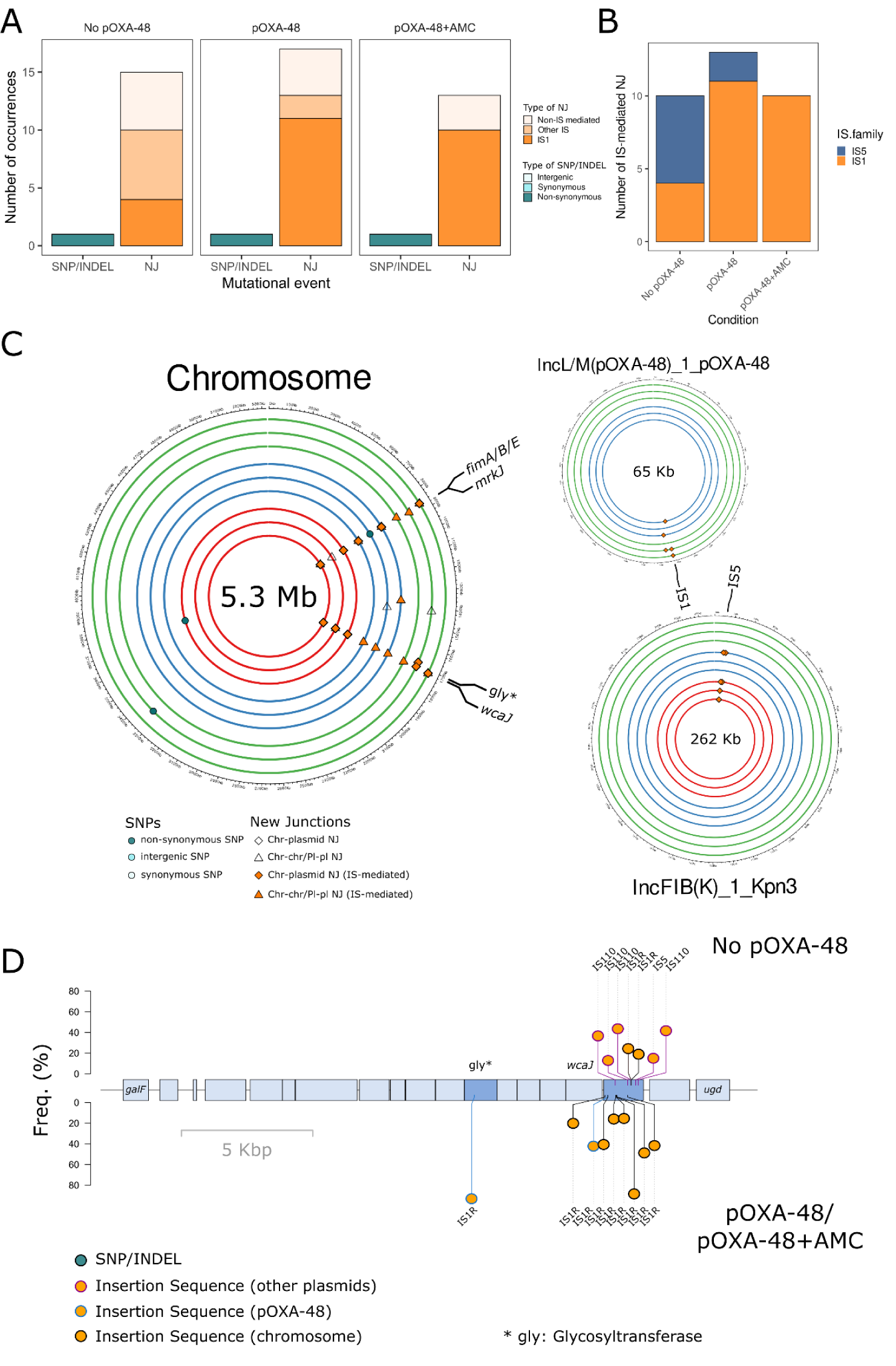
Summary of genomic changes during EE in strain H53 (*K. pneumoniae* ST487). **A)** Summary of the genetic changes in the propagated populations at day 15 classified in SNPs/INDELs or NJ for each experimental condition. Barplot colors indicate the type of event. **B)** NJ events classified by IS families per EE condition. IS1 transposition events were triggered by pOXA-48 presence during the EE. **C)** Circa plots of the H53 genome showing the location of the genomic changes. From inside to outside, circles indicate the different replicates of the evolved bacterial populations without pOXA-48 (red), carrying pOXA-48 (blue) and carrying pOXA-48 in presence of AMC (green). Parallel evolution targets are labeled in each circa plot. The different types of SNPs/INDELs are represented by dots, whereas NJ events are depicted by shapes, filled in orange in the case of IS-mediated NJs. The fimbrial operon and the capsule operon were the main targets of adaptation. **D)** Lolliplot of the capsule operon. The top part of the plot shows mutations occurred during the EE in the replicates propagated in absence of the plasmid, whereas the bottom part includes the events that happened in pOXA-48 carrying replicates both with and without AMC. Names of the genes are shown. IS elements mediated the loss of the capsule in this strain independently of pOXA-48 presence. However, the IS families which mediated the KO changed from IS5 to mainly IS1 when pOXA-48 was present during EE.

**Supplementary Figure 12.**
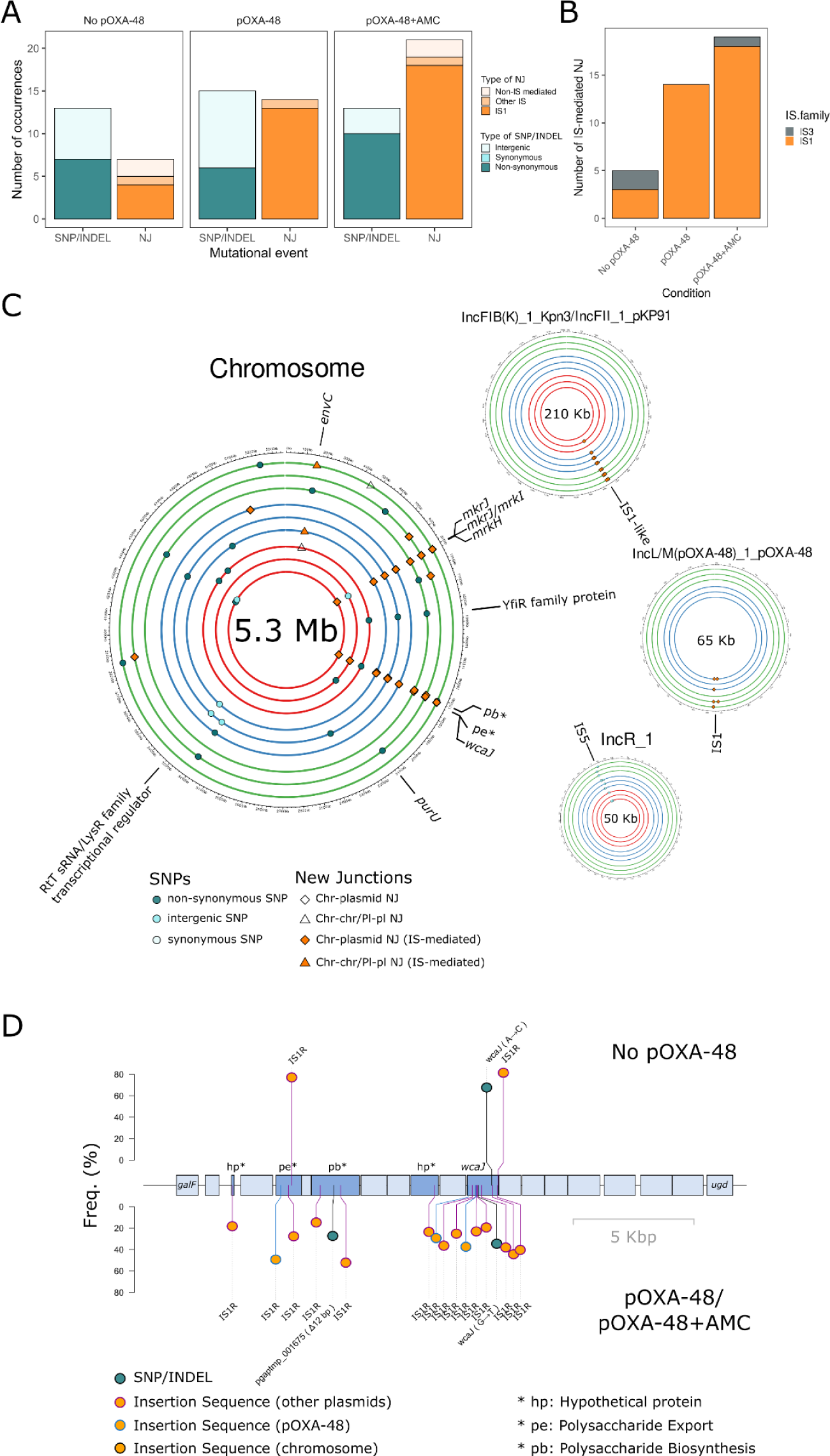
Summary of genomic changes during EE in strain K25 (*K. pneumoniae* ST11). **A)** Summary of the genetic changes in the propagated populations at day 15 classified in SNPs/INDELs or NJ for each experimental condition. Barplot colors indicate the type of event. **B)** NJ events classified by IS families per EE condition. IS1 transposition events were triggered by pOXA-48 presence during the EE. **C)** Circa plots of the K25 genome showing the location of the genomic changes. From inside to outside, circles indicate the different replicates of the evolved bacterial populations without pOXA-48 (red), carrying pOXA-48 (blue) and carrying pOXA-48 in presence of AMC (green). Parallel evolution targets are labeled in each circa plot. The different types of SNPs/INDELs are represented by dots, whereas NJ events are depicted by shapes, filled in orange in the case of IS-mediated NJs. The fimbrial operon and the capsule operon were the main targets of adaptation. **D)** Lolliplot of the capsule operon. The top part of the plot shows mutations occurred during the EE in the replicates propagated in absence of the plasmid, whereas the bottom part includes the events that happened in pOXA-48 carrying replicates both with and without AMC. Names of the genes are shown. IS elements mediated the loss of the capsule in this strain independently of pOXA-48 presence. However, the IS families which mediated the KO changed from IS5 to mainly IS1 when pOXA-48 was present during EE.

**Supplementary Figure 13.**
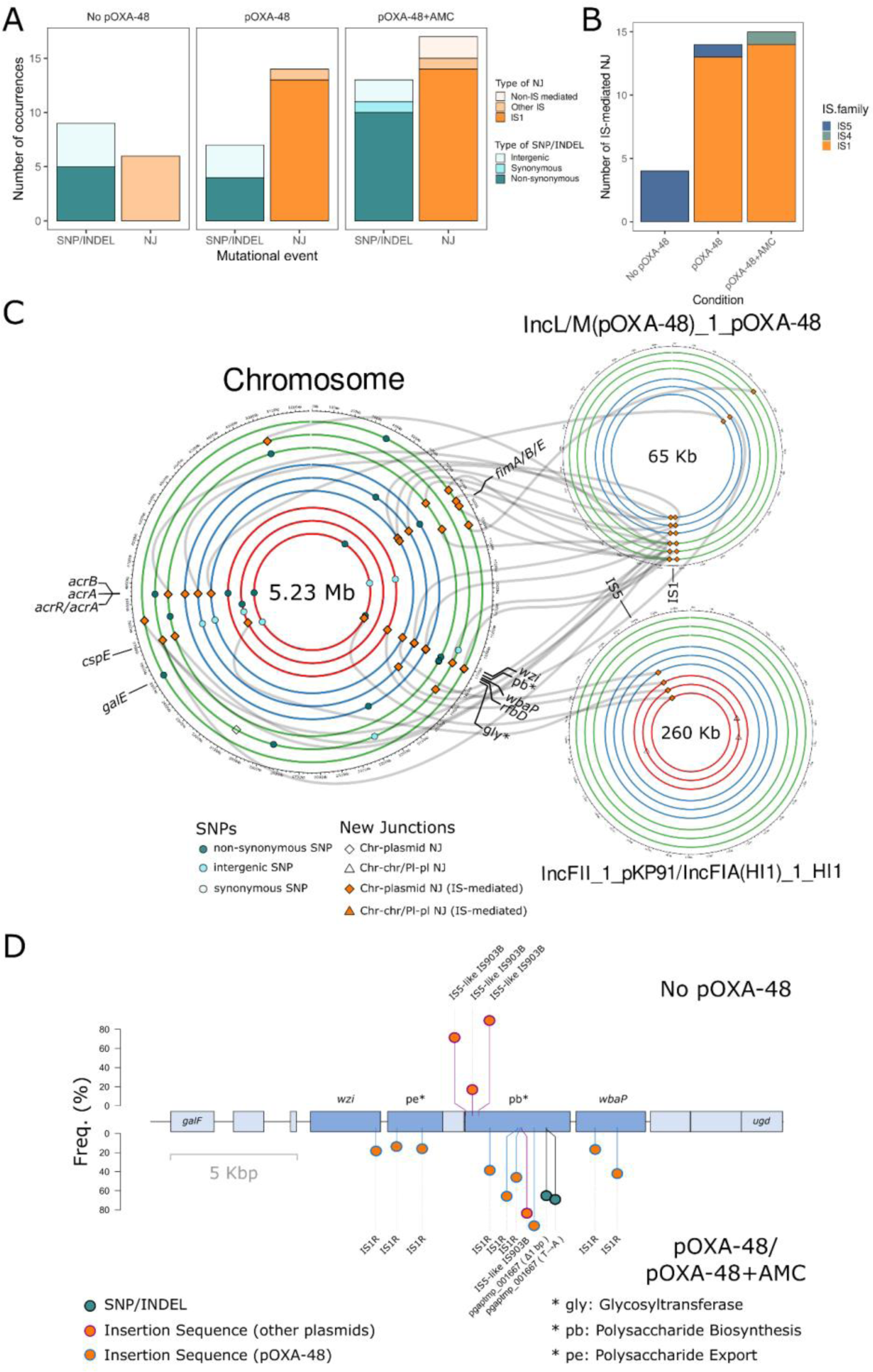
Summary of genomic changes during EE in strain K091 (*K. pneumoniae* ST1427). **A)** Summary of the genetic changes in the propagated populations at day 15 classified in SNPs/INDELs or NJ for each experimental condition. Barplot colors indicate the type of event. **B)** NJ events classified by IS families per EE condition. IS1 transposition events were triggered by pOXA-48 presence during the EE. In the absence of pOXA-48 we detected IS5 transposition events, which result almost insignificant in pOXA-48-carrying populations. **C)** Circa plots of the K091 genome showing the location of the genomic changes. From inside to outside, circles indicate the different replicates of the evolved bacterial populations without pOXA-48 (red), carrying pOXA-48 (blue) and carrying pOXA-48 in presence of AMC (green). Parallel evolution targets are labeled in each circa plot. The different types of SNPs/INDELs are represented by dots, whereas NJ events are depicted by shapes, filled in orange in the case of IS-mediated NJs. IS rearrangements which could be tracked (i.e., confirmed by genomic data, and/or by long-read sequencing) are shown as lines connecting the IS element and its target. The fimbrial operon, the capsule operon and the acrAB efflux pump genes were the main targets of adaptation. **D)** Lolliplot of the capsule operon. The top part of the plot shows mutations occurred during the EE in the replicates propagated in absence of the plasmid, whereas the bottom part includes the events that happened in pOXA-48 carrying replicates both with and without AMC. Names of the genes are shown. In absence of pOXA-48, IS5 elements from an IncF plasmid mediated the KO of the capsule, whereas mainly IS1 elements mediated it in pOXA-48-carrying populations.

**Supplementary Figure 14.**
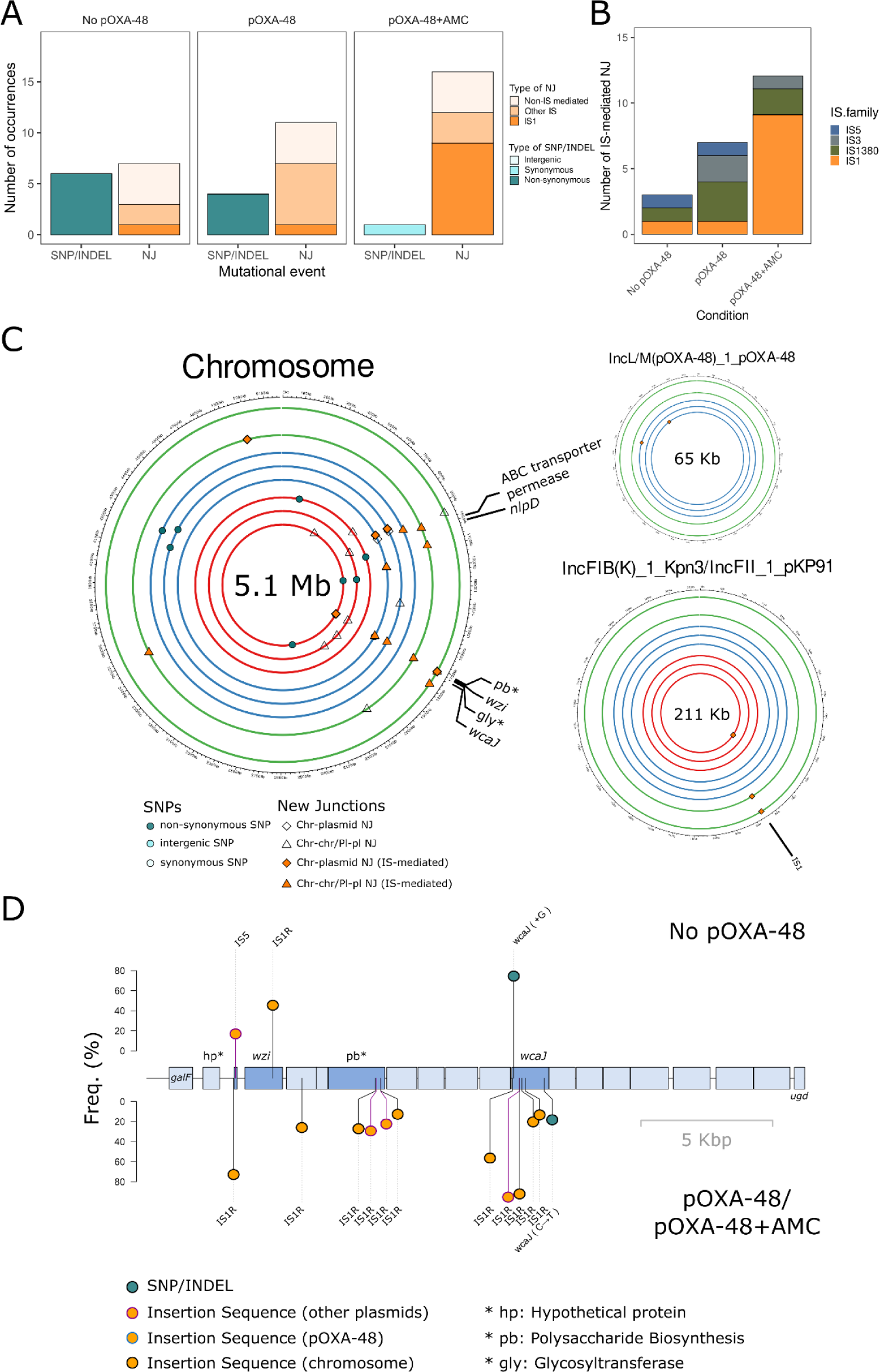
Summary of genomic changes during EE in strain K147 (*K. pneumoniae* ST11). **A)** Summary of the genetic changes in the propagated populations at day 15 classified in SNPs/INDELs or NJ for each experimental condition. Barplot colors indicate the type of event. **B)** NJ events classified by IS families per EE condition. IS1 transposition events were triggered by pOXA-48+AMC presence during the EE. **C)** Circa plots of the K147 genome showing the location of the genomic changes. From inside to outside, circles indicate the different replicates of the evolved bacterial populations without pOXA-48 (red), carrying pOXA-48 (blue) and carrying pOXA-48 in presence of AMC (green). Parallel evolution targets are labeled in each circa plot. The different types of SNPs/INDELs are represented by dots, whereas NJ events are depicted by shapes, filled in orange in the case of IS-mediated NJs. The *rpoS-nlpD* stress response operon and the capsule operon were the main targets of adaptation. We kept out of the analyses the hypermutator replicate (6_H4 mutated in *mutH*) propagated with pOXA-48 and AMC. **D)** Lolliplot of the capsule operon. The top part of the plot shows mutations occurred during the EE in the replicates propagated in absence of the plasmid, whereas the bottom part includes the events that happened in pOXA-48 carrying replicates both with and without AMC. Names of the genes are shown. In presence of pOXA-48, the number of non capsulated mutants increased due to a higher IS1 transposition rate as shown in Fig. 4E.

**Supplementary Figure 15.**
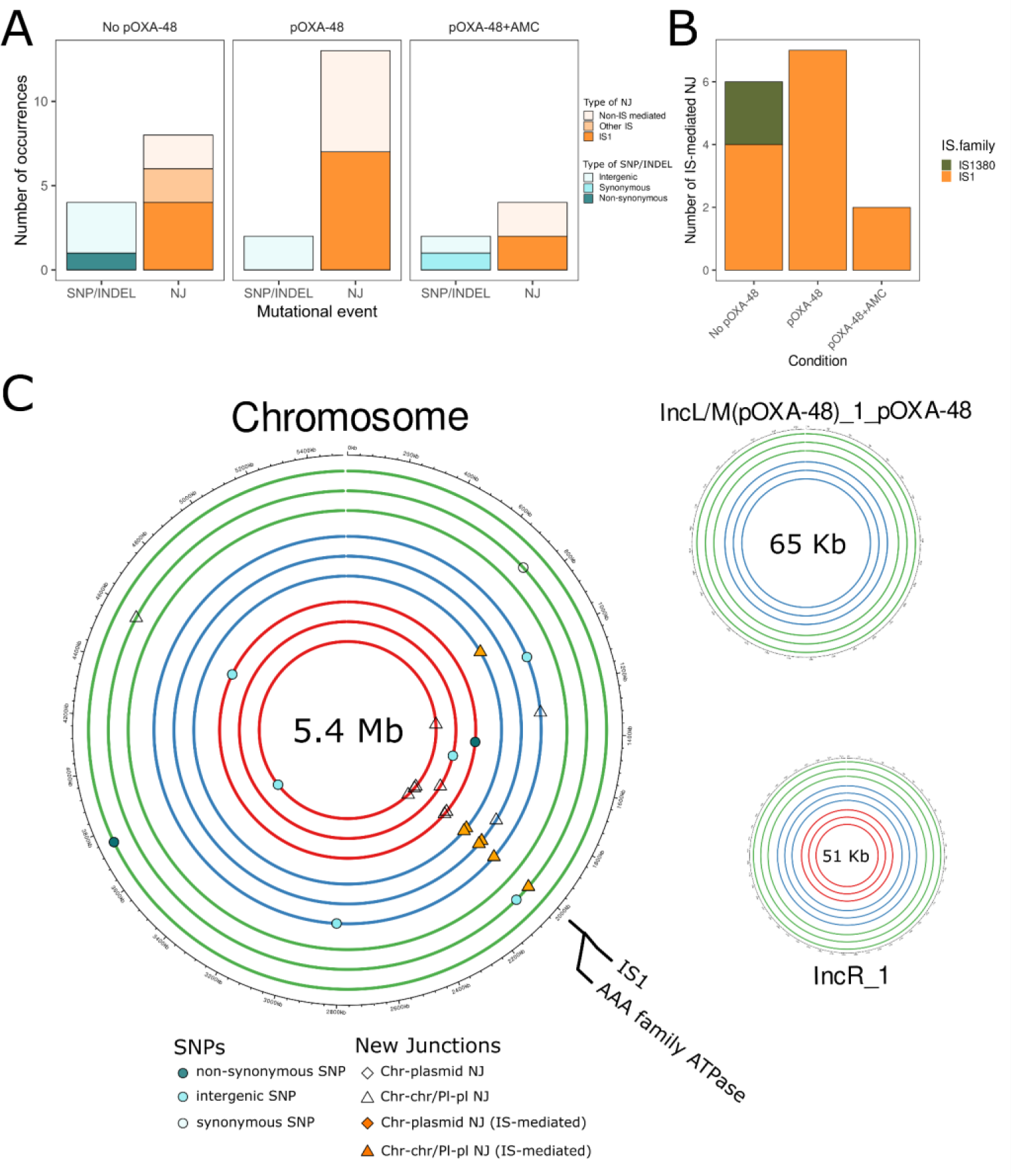
Summary of genomic changes during EE in strain K153 (*K. pneumoniae* ST11). **A)** Summary of the genetic changes in the propagated populations at day 15 classified in SNPs/INDELs or NJ for each experimental condition. Barplot colors indicate the type of event. **B)** NJ events classified by IS families per EE condition. No increase, and even a notable decrease was detected in pOXA-48 and pOXA-48+AMC populations respectively. This strain showed both the higher genomic IS1 copy number among the *K. pneumoniae* strains selected (6 copies vs 1-5; Suppl. Table 1) and its pOXA-48 variant lacked one of its IS1 elements (Suppl. Fig 2), potentially affecting its transposition activity.**C)** Circa plots of the K153 genome showing the location of the genomic changes. From inside to outside, circles indicate the different replicates of the evolved bacterial populations without pOXA-48 (red), carrying pOXA-48 (blue) and carrying pOXA-48 in presence of AMC (green). Parallel evolution targets are labeled in each circa plot. The different types of SNPs/INDELs are represented by dots, whereas NJ events are depicted by shapes, filled in orange in the case of IS-mediated NJs.

**Supplementary Figure 16.**
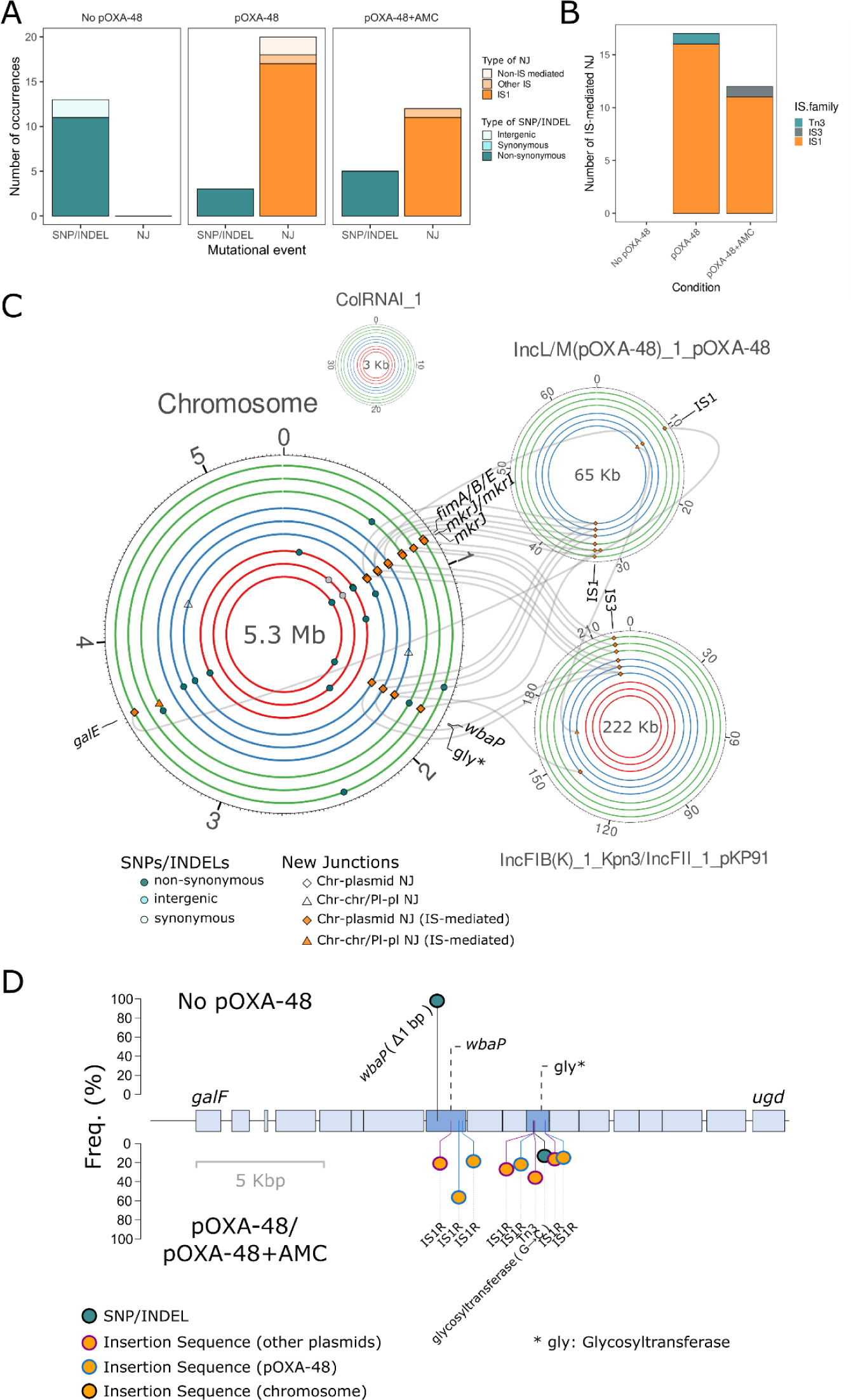
Summary of genomic changes during EE in strain K163 (*K. pneumoniae* ST307). **A)** Summary of the genetic changes in the propagated populations at day 15 classified in SNPs/INDELs or NJ for each experimental condition. Barplot colors indicate the type of event. **B)** NJ events classified by IS families per EE condition. IS1 transposition events were triggered by pOXA-48 presence during the EE. **C)** Circa plots of the K163 genome showing the location of the genomic changes. From inside to outside, circles indicate the different replicates of the evolved bacterial populations without pOXA-48 (red), carrying pOXA-48 (blue) and carrying pOXA-48 in presence of AMC (green). Parallel evolution targets are labeled in each circa plot. The different types of SNPs/INDELs are represented by dots, whereas NJ events are depicted by shapes, filled in orange in the case of IS-mediated NJs. IS rearrangements which could be tracked (i.e., confirmed by genomic data, and/or by long-read sequencing) are shown as lines connecting the IS element and its target. The fimbrial operon and the capsule operon and *galE* were the main targets of adaptation. **D)** Lolliplot of the capsule operon. The top part of the plot shows mutations occurred during the EE in the replicates propagated in absence of the plasmid, whereas the bottom part includes the events that happened in pOXA-48 carrying replicates both with and without AMC. Names of the genes are shown. The pathways of adaptation differed depending on the presence of pOXA-48, as the capsule loss is generally mediated by IS1 in plasmid carrying populations, whereas no IS1 transposition disrupting this operon could be detected in plasmid free populations.

**Supplementary Figure 17.**
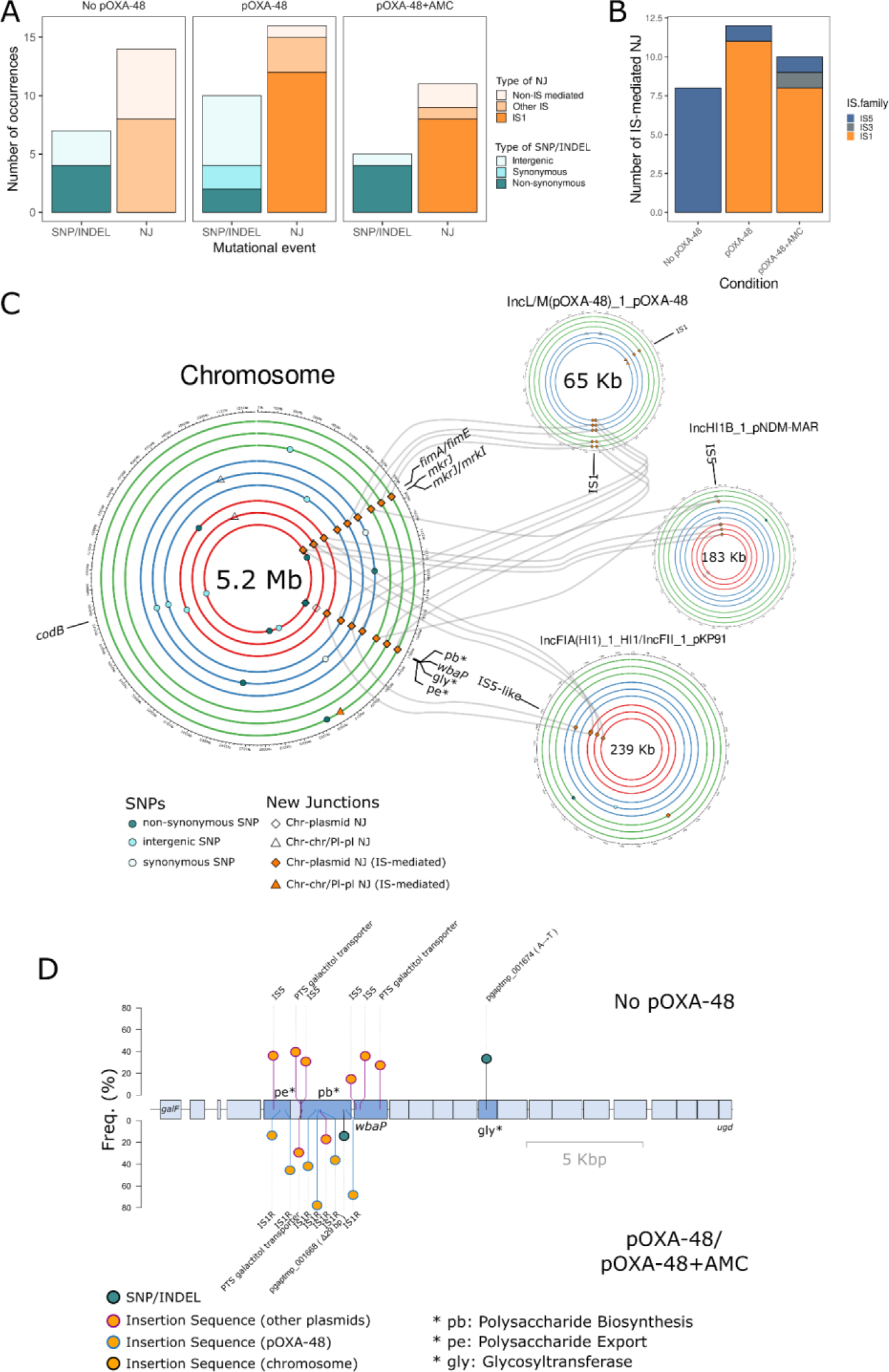
Summary of genomic changes during EE in strain K209 (*K. pneumoniae* ST1427). **A)** Summary of the genetic changes in the propagated populations at day 15 classified in SNPs/INDELs or NJ for each experimental condition. Barplot colors indicate the type of event. **B)** NJ events classified by IS families per EE condition. In the absence of pOXA-48 we detected IS5 transposition events, which result almost insignificant in pOXA-48-carrying populations. **C)** Circa plots of the K209 genome showing the location of the genomic changes. From inside to outside, circles indicate the different replicates of the evolved bacterial populations without pOXA-48 (red), carrying pOXA-48 (blue) and carrying pOXA-48 in presence of AMC (green). Parallel evolution targets are labeled in each circa plot. The different types of SNPs/INDELs are represented by dots, whereas NJ events are depicted by shapes, filled in orange in the case of IS-mediated NJs. Interestingly, Lastly, although the goal of this study was to analyze the generalized impact of pOXA-48 in the evolution of clinical enterobacteria beyond compensatory evolution, we detected interesting compensatory events in this strain, in which pOXA-48 imposed a significant cost (K209; Fig. 1B). Although at low frequencies, in 2 pOXA-48-carrying evolved populations of this strain (without AMC), we observed a partial deletion of the *bla*_OXA-48_ gene, an event we have previously described in vivo that potentially alleviates the cost of pOXA-48^55^. **D)** Lolliplot of the capsule operon. The top part of the plot shows mutations occurred during the EE in the replicates propagated in absence of the plasmid, whereas the bottom part includes the events that happened in pOXA-48 carrying replicates both with and without AMC. Names of the genes are shown. The knockout of the capsule was mediated by IS5 elements from an IncF plasmid in pOXA-48-free populations, whereas mainly IS1 mediated the knockout in presence of pOXA-48.

**Supplementary Figure 18.**
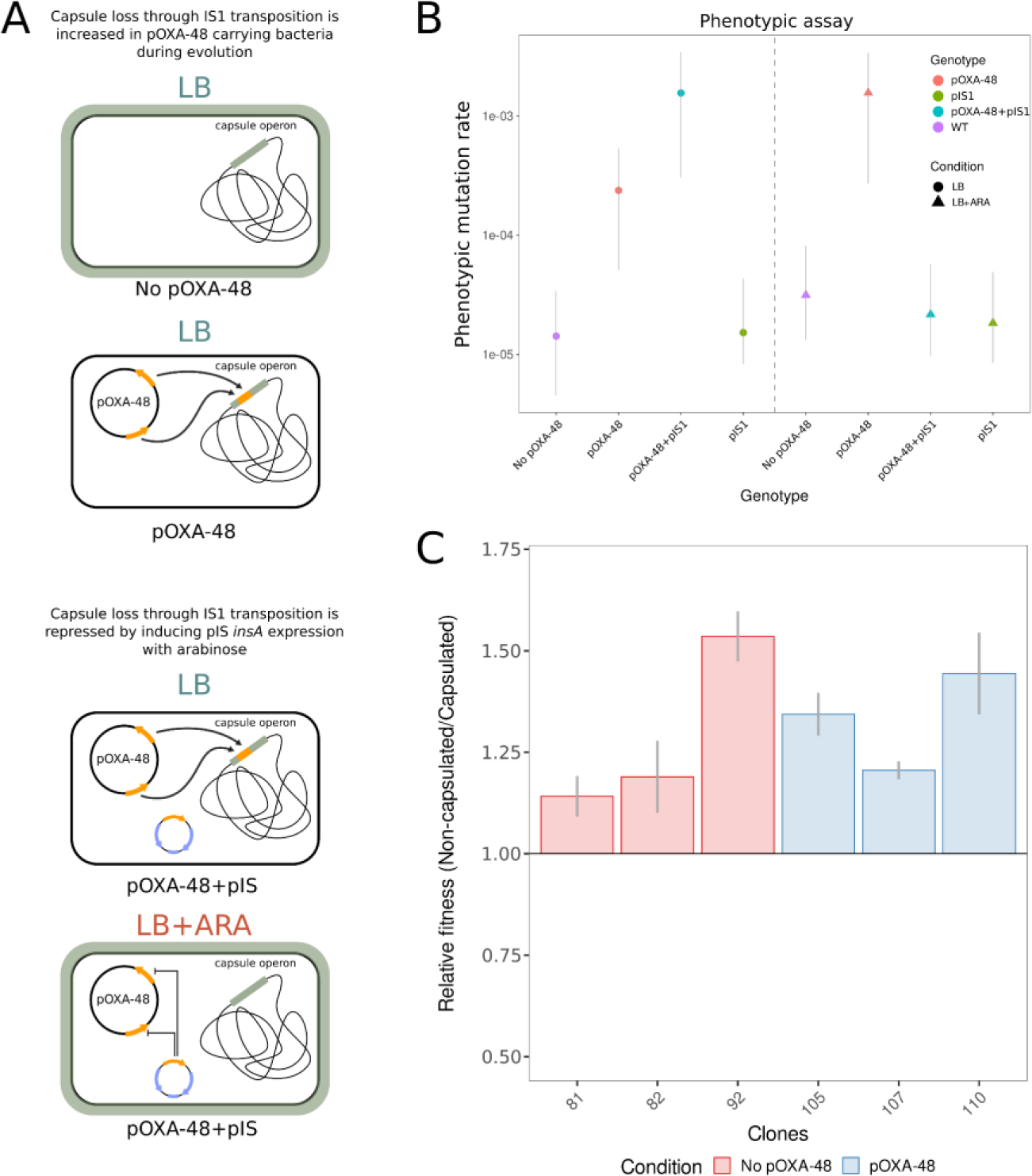
Scheme of capsule loss in the different conditions tested during the fluctuation assay of IS1 transposition and its results including all the controls. **A)** Scheme showing capsule loss in the different conditions both with and without pIS1. Our EE observations indicated that non-capsulated *K. pneumoniae* mutants appeared mainly due to IS1 insertions in the capsule operon in presence of pOXA-48. Hence, the induction of pIS1 in presence of arabinose should repress the transposition of pOXA-48 IS1 elements into the capsule operon due to increased levels of repressor (InsA). **B)** Phenotypic mutation rate of non-capsulated mutants in logarithmic scale. Left side of the plot shows all the results for the samples tested in LB, including the different phenotypes with 1, 2 or none of the plasmids (pOXA-48; pIS1), while the right plot shows the results of the fluctuation assay in presence of arabinose (LB+ARA). We could detect a clear repression of non-capsulated phenotype in pOXA-48+pIS1 samples in presence of arabinose, reaching levels very similar to those of pOXA-48-free samples. Error bars represent 95% confidence intervals. **C)** Relative fitness levels of non-capsulated vs. capsulated bacteria without and with pOXA-48 resulting from competition assays (n = 3). We selected clones with mutations in different genes along the capsule operon: pe (81 [stop codon], 105 [IS1 inactivation]), pb (82 [IS1 inactivation]), and *wcaJ* (92 [IS1 inactivation], 107 [IS1 inactivation] and 110 [stop codon]). Increase in relative fitness ranged from 14% to 53% in non-capsulated mutants, both in the presence and absence of pOXA-48.

**Supplementary Figure 19.**
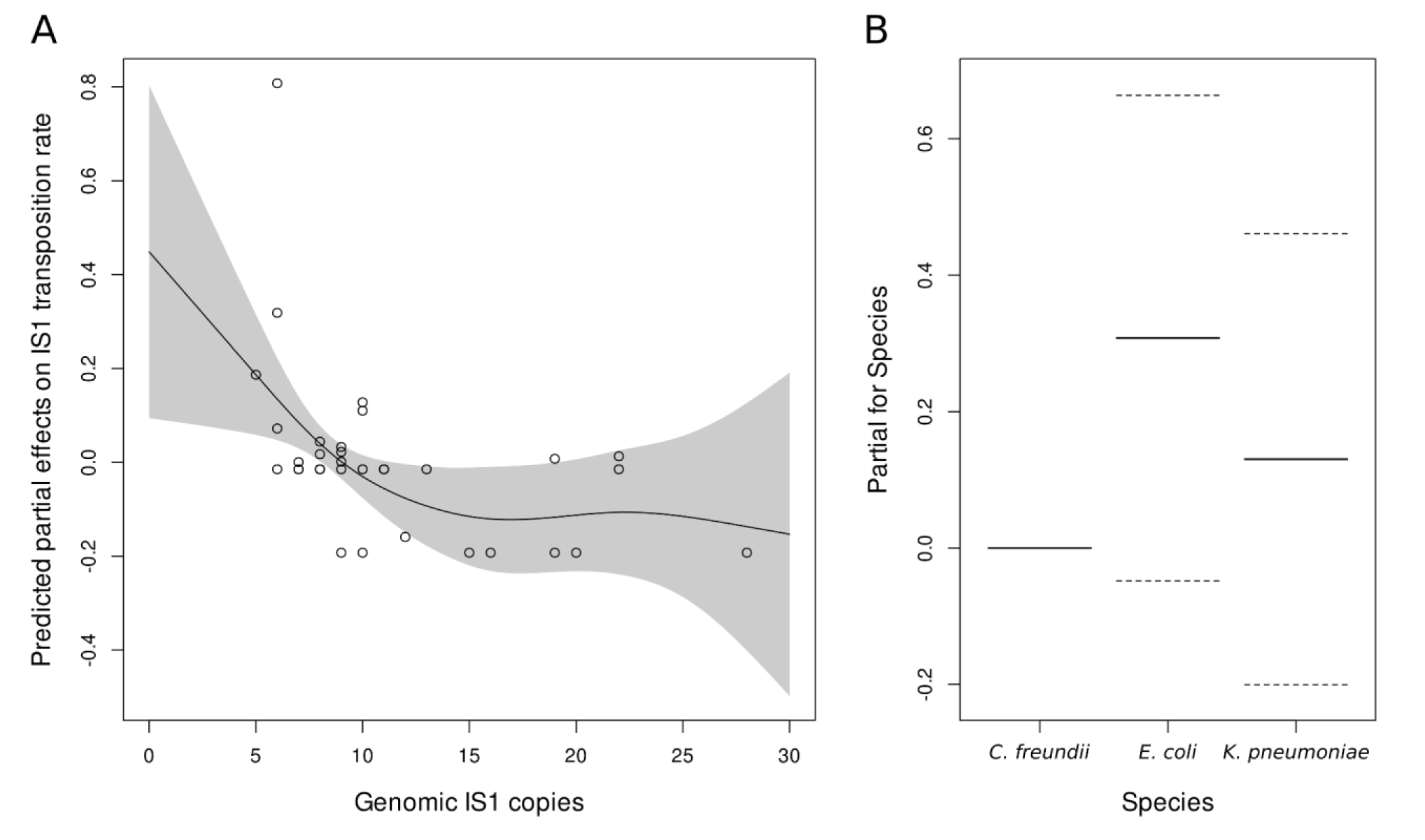
GAM model results. **A)** Predicted partial effects of genomic IS1 copies in the transposition rate of *in vivo* lineages analyzed through a Generalized Additive Model (GAM). The smooth function (number of basis functions: k = 6) shows a significant decrease in transposition rate (p = 0.032; F = 3.28; R-squared = 0.203; Deviance explained = 29.7%) as genomic IS1 copies increase in the strains, supporting our previous observations. Grey shade indicates the 95% confidence interval area. **B)** Predicted partial effects for the different species (parametric coefficients) included in the GAM. No significant differences were reported between species (Intercept t = -0.704, p = 0.4864; *E. coli* t = 1.73, p = 0.0929; *K. pneumoniae* t = 0.1654, p = 0.4366). Solid lines indicate the mean partial effect and dashed lines indicate the 95% confidence interval for each species.

**Supplementary Figure 20.**
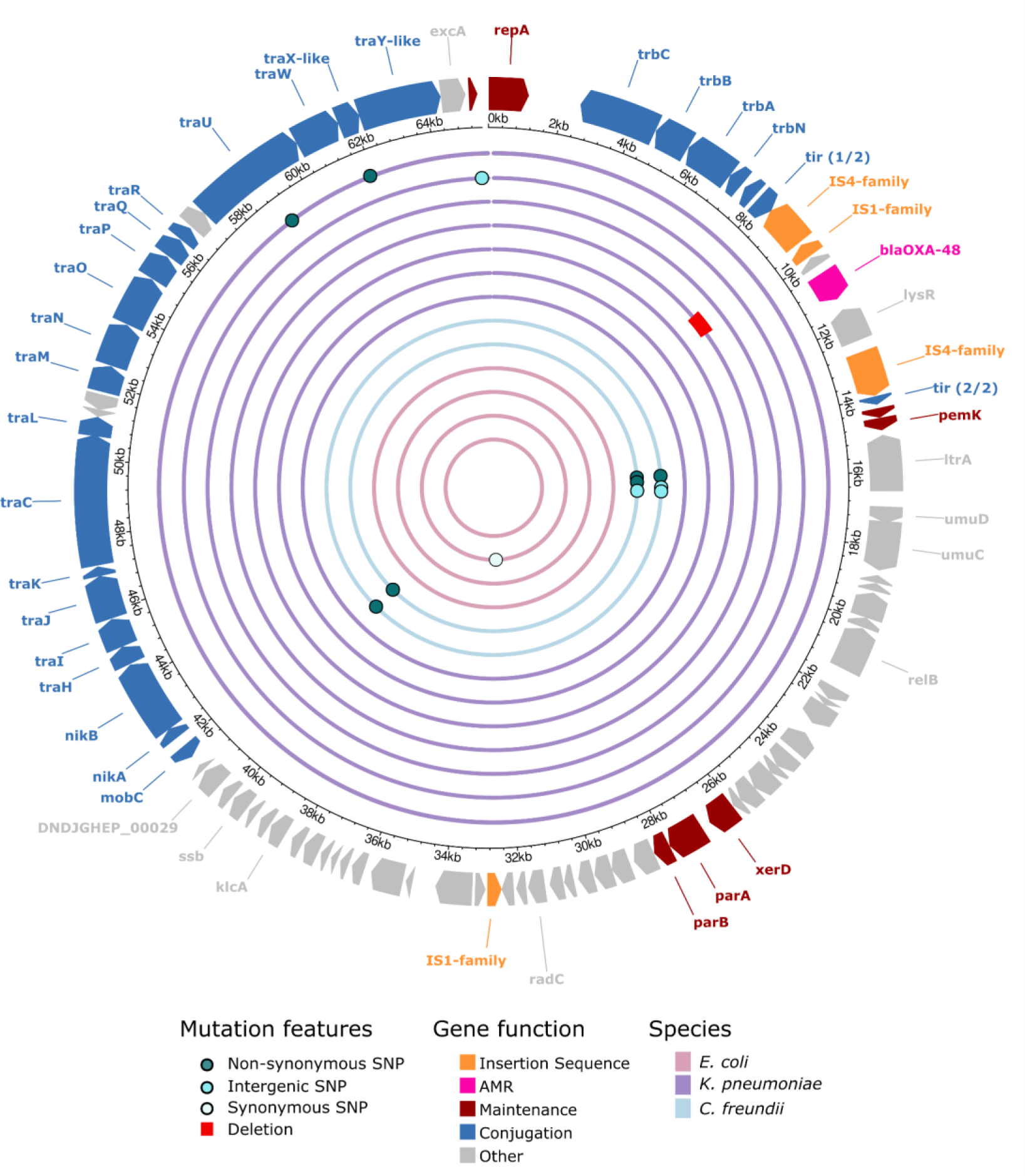
pOXA-48 variants detected in the selected strains: Scheme of pOXA-48 variants in the selected strains for this work as compared against the most common variant of pOXA-48 (K8). Each concentric circle represents one plasmid. From the inner to the outer, the strains are: C021, C286, C309, C324, CF12, CF13, H53, K091, K147, K153, K163, K209, K25. Most mutations found were accumulated in *ltrA*, a mobile element of the plasmid, in both *C. freundii* strains, as well as a non-synonymous SNP in a hypothetical protein of unknown function. For *E. coli*, only one strain showed a synonymous mutation (C286), whereas for *K. pneumoniae* strains, three showed differences compared with the K8 variant. Interestingly, K153 showed a deletion in the IS1 element around the 10 kbp, which could potentially affect transposition activity. This strain did not show plasmid-mediated IS1 transposition. K209 showed an intergenic SNP in the region between *repC* and *repA*, potentially increasing pOXA-48 PCN. K25 showed two SNPs in conjugation related genes.

**Supplementary Figure 21.**
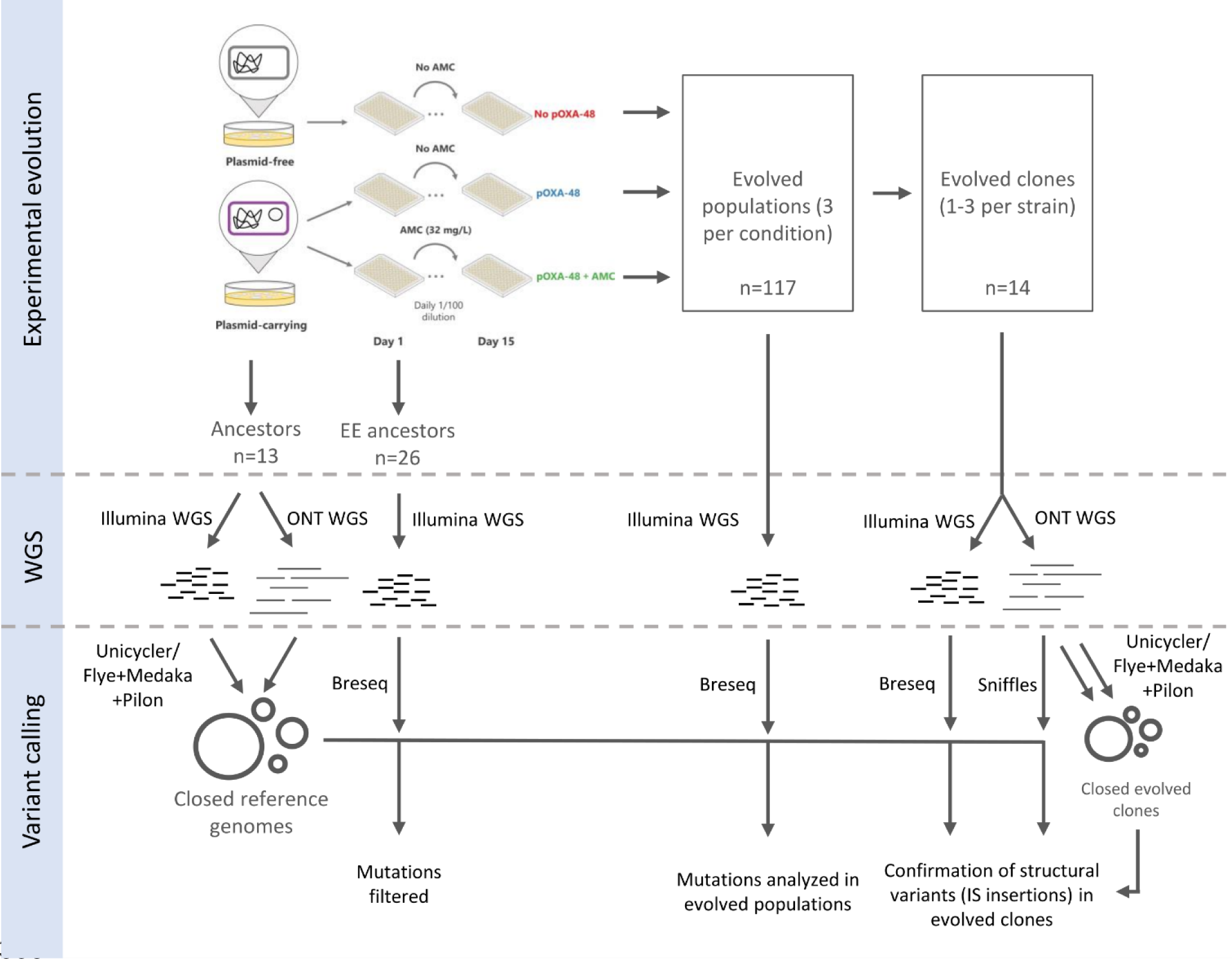
Schematic representation of EE+WGS workflow. This scheme summarizes the main technologies used to sequence each of the samples analyzed during the experimental evolution (Illumina = short-read; Oxford Nanopore Technologies (ONT) = long-read). We combined both short and long read sequencing for getting hybrid closed assemblies of both the ancestors of each strain and of the evolved clones isolated from the populations at day 15. We mainly used short read WGS to filter out possible mutations at day 1 (as a control for the EE), as well as for analyzing the mutations present in the evolved populations and isolated clones at day 15. We used the hybrid assemblies of the ancestors as the reference for the variant calling both populations and clones. From the 117 evolved populations, we discarded 4: a contaminated replicate of *C. freundii* without pOXA-48, a contaminated replicate of *E. coli* due to low mapping % when performing the variant calling, and the two hypermutator strains indicated in the main text. We used breseq to perform the variant calling with short-read data. To confirm the results reported by breseq we complemented the variant calling of the clonal samples using long-read data and the Sniffles software (specifically developed to detect Structural Variants). Finally, we also supported the variant calling results with the closed genomes of the evolved clones. We indicate the sequencing technology or the software used for the hybrid assembly/variant calling next to each arrow. We show input samples and analysis results inside rectangles.

